# A transcriptional continuum from clinically normal to lesional skin: Single-cell trajectory analysis of psoriasis

**DOI:** 10.64898/2026.09.15.751704

**Authors:** Gülnur Uzun, Deniz Konak, Hilal Kazan, Pınar Pir

**Affiliations:** Gebze Technical University, Department of Bioengineering, Kocaeli, Türkiye; Antalya Bilim University, Department of Computer Engineering, Antalya, Türkiye

**Keywords:** Psoriasis, single-cell RNA sequencing, pseudotime trajectory, transcriptional continuum, keratinocytes, endothelial cells

## Abstract

Psoriasis is a chronic inflammatory skin disease characterized by immune dysregulation and complex cellular interactions involving keratinocytes (KCs), T cells, and endothelial cells. Single-cell studies have mapped the cellular diversity of psoriatic plaques, yet most analyses treat skin as either healthy or diseased and therefore overlook the graded changes that precede a visible lesion. We asked whether psoriasis instead progresses along a transcriptional continuum and whether clinically normal skin from patients already carries a measurable disease signature.

Using single-cell RNA sequencing (scRNA-seq) of healthy skin (NS), clinically normal skin adjacent to lesions (PN), and lesional skin (PP), we combined two complementary designs: a complete set spanning all three states and a within-patient paired set of matched PN and PP skin biopsies that removes inter-individual variability. Trajectory analysis using a novel pseudotime approach reveals transcriptional changes within cell populations and dynamic cellular states underlying psoriatic pathology. To distinguish lesion-driven effects from disease-associated transcriptional changes, we applied a differential expression strategy comparing PP and PN psoriatic tissue with NS and integrating these contrasts. This approach enabled the identification of disease-specific gene signatures while minimizing lesion-specific bias.

As a result, we identified a coherent, cell-type-resolved continuum of transcriptional states in which endothelial cells, IL20+ keratinocytes (KCs: IL20), and KGF+ keratinocytes (KCs: KGF) undergo trajectory-dependent state transitions. These trajectories converge on features including epidermal thickening, barrier dysfunction, and plaque maintenance. Endothelial cells transition from an inflamed, angiogenic state to a mature, barrier-stabilized phenotype, while KCs: IL20 and KCs: KGF shift from PN states toward hyperproliferative, stress-adapted lesional phenotypes. These findings reframe psoriasis as a dynamic, multicellular continuum and identify candidate temporal windows for early intervention.

**Key Points:**

- Psoriatic skin states form a continuum, with a clinically normal skin from patients occupying an intermediate position between healthy and lesional skin rather than aligning with either extreme.
- A within-patient paired design of matched non-lesional and lesional biopsies isolates lesion-associated changes from inter-individual variability, while a complete three-state design captures the full healthy-to-lesional gradient.
- Pseudotime trajectories resolve coordinated reprogramming of endothelial cells and two KC subsets, IL-20+ and KGF+, each transitioning from homeostatic toward hyperproliferative, stress-adapted states.
- A node-anchored differential expression strategy links transcriptional transitions directly to shifts in tissue identity along trajectory branches.
- Intermediate cells retain protective, homeostatic features, suggesting a plastic state that may delay full psoriatic transformation and define temporal windows for therapeutic intervention.

## 1. INTRODUCTION

Psoriasis is a chronic inflammatory skin disorder marked by sharply demarcated red plaques, scaling, and itch, and is driven principally by dysregulation of the immune system. Although not life-threatening, the disease imposes a substantial burden on physical health and psychosocial well-being [1], and its reported global prevalence ranges from 0.09% to 11.43%, placing psoriasis among the most common chronic skin diseases [2]. The condition is now understood as an immune-mediated disease that emerges in genetically susceptible individuals, in whom environmental and antigenic triggers, including infection, certain medications, skin injury, alcohol, smoking, and psychological stress, initiate and amplify cutaneous inflammation [1].

Single-cell RNA sequencing (scRNA-seq) has transformed the research on cellular heterogeneity, resolving disease related cell populations in psoriasis and psoriatic arthritis, supporting targeted therapy and earlier diagnosis, and revealing previously unrecognized subtypes that participate in disease progression. Psoriatic skin harbors diverse subsets of T cells, keratinocytes (KCs), dendritic cells (DCs), and other immune populations. Keratinocytes, the predominant epidermal cell type, serve not only as structural components of the skin barrier but also as active participants in the onset and persistence of disease. As sentinels of innate immunity, stressed keratinocytes release self-DNA, self-RNA, and antimicrobial peptides that form complexes capable of activating Toll-like receptors on plasmacytoid DCs and myeloid DCs; the resulting type I interferon (IFN) production and proinflammatory cytokine secretion trigger psoriatic inflammation [3]. KCs also sense nucleic acids through TLR3 and MAVS, further raising expression of TNF, IL-6, and IFN. Keratinocytes, therefore, act as both effectors and regulators of psoriatic pathology and shape the response to treatment. Biologics directed against the TNF, IL-23, and IL-17 axes effectively suppress the inflammation and normalize keratinocyte proliferation, yet relapse and phenotypic switching remain clinical challenges [3].

Among the cytokines that organize the response, interleukin-20 (IL-20), a proinflammatory member of the IL-10 family, occupies a central position. Unlike IL-10, IL-20 promotes inflammation and is markedly upregulated in lesional KCs, dermal capillaries, and epidermal leukocytes [4]. Signaling through IL-20R1/IL-20R2 and IL-22R1/IL-20R2 complexes activates the JAK–STAT3 and MAPK pathways, driving KC hyperproliferation, release of cytokines, chemokines, and antimicrobial peptides, and increased angiogenesis, which together amplify local inflammation [5]. IL-20+ keratinocytes form a distinct cluster of cells, positioning IL-20 as a key link between immune infiltration, keratinocyte behavior, and local inflammation and as a candidate biomarker and therapeutic target.

A second KC program centers on keratinocyte growth factor (KGF, also known as fibroblast growth factor 7, *FGF7*). Among the FGF family, KGF acts selectively on epithelial cells, exerting strong mitogenic effects on KCs while sparing fibroblasts and endothelial cells. Binding to the epithelial receptor FGFR2b activates the MAPK/ERK and PI3K/AKT pathways, supporting epithelial proliferation, differentiation, and tissue repair, and thereby underpinning skin homeostasis and wound healing. Aberrant KGF signaling has been linked to skin disorders including psoriasis, where excessive keratinocyte growth contributes to disease [6].

A recent integrative transcriptomic analysis of psoriasis identified immune-pathway enrichment among disease-associated genes, including cytokine-cytokine receptor interaction and IL-17 signaling [7]. In the same study, single-cell analysis described extensive intercellular communication among KCs, DCs, monocytes, and T cells. Trajectory analysis further identified distinct states enriched for IL1B-expressing KCs and T cells versus other states mainly comprising T cells together with IFNG-expressing KCs. The analysis further highlighted the presence of hub genes associated with cellular composition and functional status in the psoriatic immune microenvironment [7]. The single-cell dataset analyzed here (GSE173706) has been examined in two prior studies. Merleev et al. [4] generated these profiles as part of the study identifying *PCSK9* as a psoriasis-susceptibility locus, showing that KCs are the principal cutaneous source of *PCSK9* and that *PCSK9* and *IL36G* are present across epidermal layers. Ma et al. [8] subsequently combined single-cell and spatial transcriptomics to dissect these skin compartments, resolving KC and fibroblast heterogeneity, an IL-36-dependent amplification of IL-17A and TNF responses within the supraspinous epidermis layer, and disease-associated ligand–receptor interactions. They further ordered KCs along pseudotime, characterizing from basal to supraspinous differentiation programs in healthy and lesional psoriasis skin. These studies establish that psoriatic skin comprises distinguishable transcriptional states and begin to characterize the position of clinically normal patient skin relative to healthy and lesional tissue. However, because they largely treat disease states as discrete categories, healthy versus diseased, and therefore cannot separate the low-level, subclinical activation present in clinically normal skin surrounding lesions from the changes restricted to established plaques. This distinction is further obscured by inter-individual variability. Because PN and PP can be sampled from the same individual, matched PN–PP comparisons offer a route to isolate lesion-associated signatures while characterizing the subclinical state of skin that appears healthy.

Motivated by these observations, we hypothesized that psoriatic skin states form a continuum rather than a set of discrete categories, spanning healthy skin, clinically normal patient skin, and overt lesions. To test this, we analyzed publicly available scRNA-seq data [4] from NS, PN, and PP in two modules: a complete set capturing differences between healthy controls and patients across all three states, and a paired set of matched PN and PP samples that controls for inter-individual variability within patients. Using a single-cell framework that integrates batch correction, cell type annotation, differential expression, and pseudotime trajectory inference, we identified disease-associated gene signatures and cell type specific programs of progression. Pseudotime analysis, in particular, exposed dynamic, tissue-dependent transcriptional changes along coherent cellular lineages, pinpointing the populations and pathways that accompany early inflammatory change. Together, these analyses provide a single-cell view of psoriatic transcriptional dynamics and nominate molecular markers of disease development.

## 2. METHODS

The scRNA-seq data (GEO accession GSE173706) were obtained from the NCBI database and comprise skin biopsies across three states: healthy normal skin (NS), clinically normal peripheral skin from patients (PN), and lesional psoriatic skin (PP) [4]. Of the 33 deposited biopsies, 30 were included in the analysis: 8 normal skin (NS), 11 non-lesional psoriatic skin (PN), and 11 lesional psoriatic skin (PP) samples. The remaining three biopsies were additional samples obtained from the same patients and corresponding tissue states already represented in the dataset. Therefore, only one biopsy per patient was retained to avoid treating non-independent samples as separate biological replicates. Two overlapping analysis sets were defined. The complete set included all three tissue states (8 NS, 11 PN, and 11 PP) and enabled comparisons across the full continuum from healthy to lesional skin. The paired set, a subset of the complete set, included 11 matched PN–PP sample pairs from the same patients and was used to identify lesion-associated changes while minimizing the effects of inter-individual variability.

### 2.1. Pre-processing of the Data

Raw gene expression matrices were downloaded in CSV format and processed with Seurat v5 [9] in R [10]. Each of the 30 samples was read individually, and gene identifiers were mapped from Ensembl IDs to gene symbols using org.Hs.eg.db [11]. Genes without valid symbols were excluded, and duplicated gene symbols were aggregated by computing the mean expression.

Each sample was converted into a Seurat object by retaining cells with more than 500 detected features and genes detected in at least three cells. Metadata labels were assigned according to sample origin, and all samples were merged into a single Seurat object for joint analysis. Quality-control metrics, including the percentage of mitochondrial gene expression, were calculated, and cells with less than 10% mitochondrial gene expression were retained. The data were then normalized using the LogNormalize method, and the top 2,000 highly variable genes were identified using the variance-stabilizing transformation (VST) method.

#### Dimensionality reduction, batch correction, and clustering

Principal component analysis (PCA) was applied with state specific settings: the first 15 principal components (PCs) were used for the complete set, and the first 10 PCs for the paired set. More PCs were used for the complete set since it contains more samples and cells, and therefore greater overall variance. Cells were embedded and visualized with UMAP [12], and unsupervised Louvain clustering was performed at resolution 0.5. Doublets were identified and removed with scDblFinder [13]. To account for batch effects and inter-sample variability, each sample was treated as a separate batch and integrated using Harmony [14]. UMAP visualization and clustering were then performed using the Harmony-corrected embeddings.

#### Cell type annotation

Cells were annotated with SingleR [15] against the Human Primary Cell Atlas reference from the celldex package, using both main (label.main) and fine (label.fine) labels. The Harmony-integrated object served as the test dataset, and predicted labels were validated against established cell type markers from the literature before being added to the object metadata.

#### Differential expression and enrichment

Differentially expressed genes (DEGs) were identified with Seurat’s *FindMarkers()* function using the Wilcoxon rank-sum test, with significance defined by an adjusted p-value below 0.05, an absolute log2 fold change above 1, and detection in at least 10% of cells. DEGs were first computed for two designs: the paired comparison of PP against matched PN and a three-state comparison incorporating NS from healthy donors. To define candidate disease markers, the DEG lists (PP vs PN, PP vs NS, and PN vs NS) were further partitioned into up- and downregulated genes under the same thresholds, and the resulting gene sets were subjected to functional enrichment with Metascape [16].

#### Pseudotime trajectory inference

Trajectories were inferred with Monocle3 [17,18]. The Seurat object was split by cell type, and pseudotime was computed separately for IL-20-high keratinocytes (KCs: IL20), KGF-high KCs (KCs: KGF), neurons, and endothelial cells. For each subset, filtered raw counts were supplied as a cell_data_set, with Monocle3 performing preprocessing. Low-dimensional manifolds were computed by UMAP, cells were grouped by community detection to identify coherent clusters and disconnected partitions, and the global graph structure was inferred with *learn_graph*, which fits a principal graph to the manifold by reversed graph embedding to capture putative paths and branch points. Cells were ordered with *order_cells* in unsupervised mode, in which root and terminal nodes are determined automatically rather than specified by the user. Pseudotime values were derived as normalized shortest-path distances from the root along the graph, and trajectories, branch points, and partitions were visualized with *plot_cells*, with cells colored by pseudotime, patient, and tissue. The analysis focused on tissue-associated structure within each cell type, prioritizing trajectory branches containing cells from more than one tissue. Because the data are cross-sectional, pseudotime is reported throughout as an ordering of transcriptional states along the inferred graph and not as a measure of elapsed time or of disease severity.

#### Node-anchored differential expression along trajectories

To compare transcriptional programs between the most divergent populations along a trajectory, we developed a node-anchored differential expression strategy using the *choose_cells* function in Monocle3. Within a branch, the most distinct nodes were identified, and the cells clustered at the edge of each node were selected, while intermediate cells were excluded. The rationale is that cells positioned farthest apart along a trajectory differ most in transcriptional state, hence, contrasting node-anchored populations sharpens the detected signal. Selected cells were stripped of their Monocle3-derived values and merged into a new Seurat object that preserved each cell’s original trajectory and region of origin. DEGs between tissue regions were identified using the same thresholds: an adjusted *p*-value < 0.05, an absolute log2 fold change > 1, and expression in at least 10% of cells. This analysis enabled a focused comparison of regional gene-expression differences within a shared lineage.

#### Genes correlated with pseudotime

Genes whose expression is correlated with pseudotime were also examined. Genes with an absolute Spearman correlation coefficient above 0.1 and an adjusted p-value below 0.01 were considered significantly correlated and retained. Because these genes vary continuously along the inferred trajectory, their enriched Gene Ontology (GO) terms were assessed with Metascape against the background of genes detected in the dataset [16].

#### Disease-specific signatures

To identify the genes that can be considered as disease-specific signatures for psoriasis, we have performed three pairwise DEG analyses:

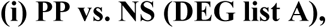

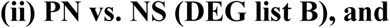

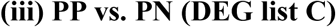

We further filtered those gene lists to be able to perform a differential gene expression assessment strategy to dissect lesion-specific (List C), non-lesional (List B), and disease-associated (List E) transcriptional changes in psoriasis. Based on DEG lists A-B-C, we subset two additional gene sets, D and E:

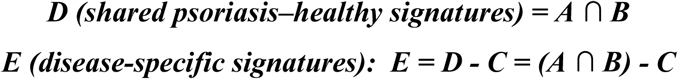

In other words, gene set E *(disease-specific signatures)* represents genes that are consistently altered in both lesional and non-lesional psoriasis relative to healthy tissue, while disregarding the genes that are present in the lesional vs. non-lesional comparison. Thus, E defines a lesion-independent, disease-associated transcriptional component that differentiates psoriasis patients from healthy control individuals, independent of local lesion formation, highlighting transcriptional differences associated with psoriasis status compared with healthy tissue.

## Results and Discussion

### Cell composition is balanced across tissue states

Following Harmony integration, major cell populations contained contributions from multiple samples, and no single sample consistently dominated the principal clusters (Fig 1). Cell-type proportions nevertheless varied between samples and tissue states, as expected in heterogeneous skin biopsies (Fig 2). Such variation may reflect biological differences between donors and tissue states as well as technical factors, and the present figures cannot distinguish between these; we therefore treat the integration as sufficient for joint visualization and clustering rather than as evidence that batch effects are absent. Cells within each group were pooled for between-group expression comparisons.

**Fig 1.**
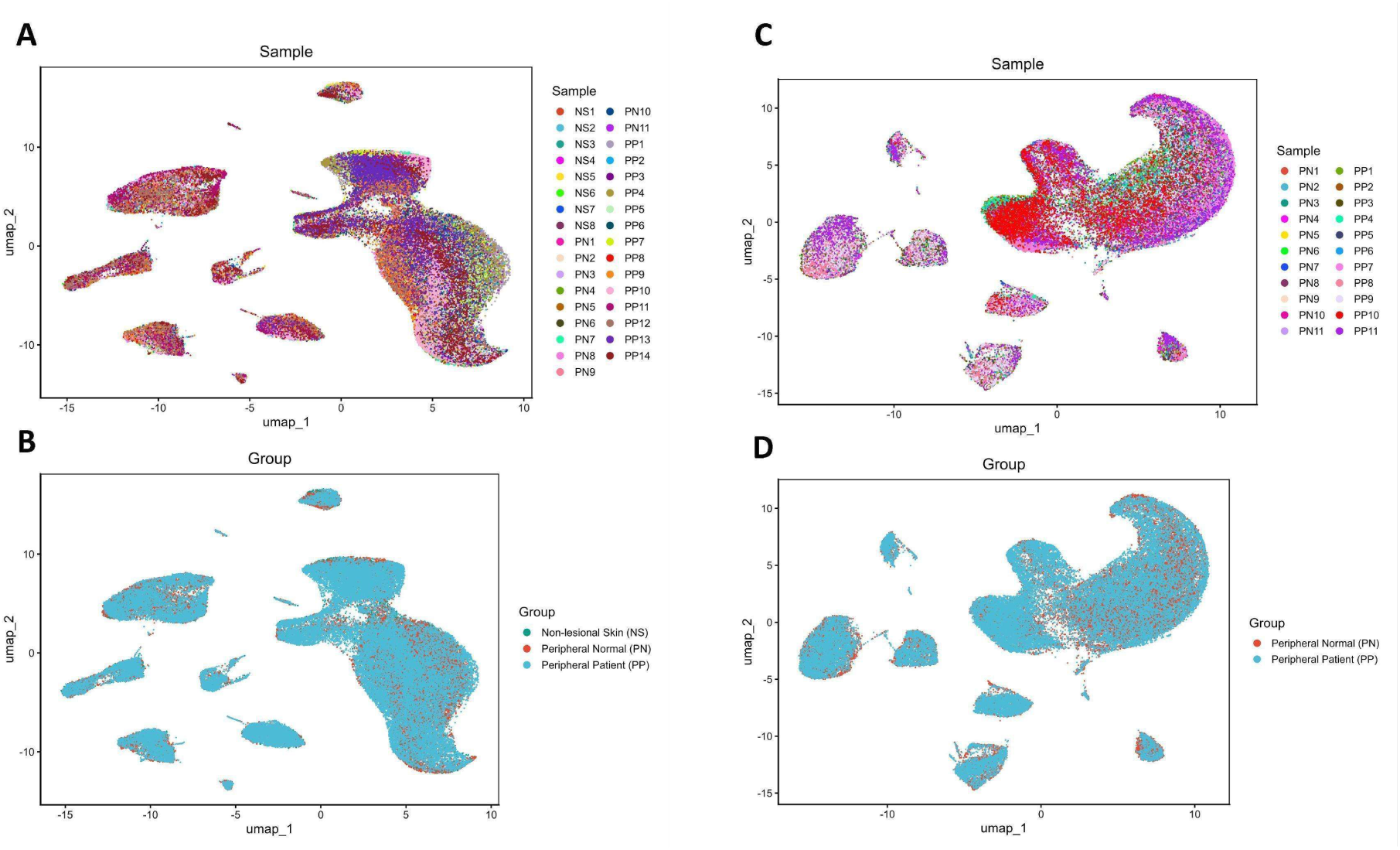
Tissue and sample distributions for both objects. (A-B) Cells color-coded for samples and tissue groups in the complete set. (C-D) Cells color-coded for samples and tissue groups in the paired set. Tissue groups are defined as NS, healthy-control skin; PN, clinically normal (non-lesional) psoriatic skin; PP, lesional psoriatic skin.

**Fig 2.**
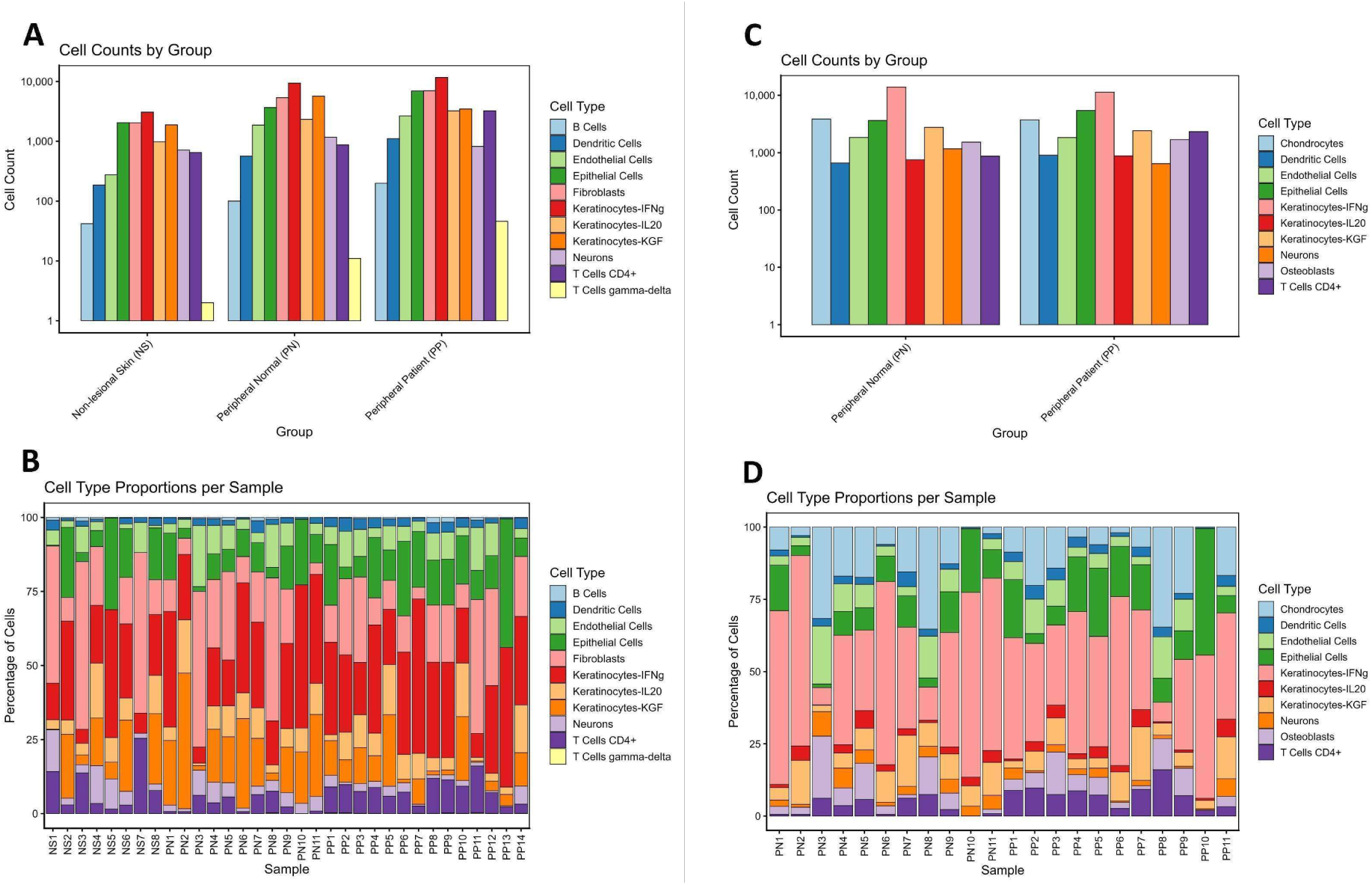
Cell count statistics. (A) Cell counts by group for the complete set. (B) Cell type proportions per sample for the complete set. (C) Cell counts by group for the paired set. (D) Cell type proportions per sample for the paired set. NS, healthy-control skin; PN, non-lesional psoriatic skin; PP, lesional psoriatic skin.

### Lesional skin diverges sharply from healthy skin

Gene expression patterns in PP and NS differed markedly. Relative to healthy skin, lesions showed heightened interferon and cytokine signaling, activation of both innate and adaptive immunity, and wound-associated tissue dynamics, alongside broad suppression of developmental, circadian, and epigenetic control. Upregulated genes were dominated by cytokine signaling, interferon-α/β signaling, interferon-stimulated gene (ISG) responses, and response to interferon-β, indicating a sustained, interferon-biased transcriptional state. Innate and adaptive pathways, including neutrophil degranulation, response to bacterium, metal sequestration by antimicrobial proteins, humoral immunity, and MHC assembly, pointed to myeloid activation, strong antimicrobial activity, and active antigen presentation. Regulatory terms such as cellular response to cytokine stimulus and regulation of inflammatory and apoptotic signaling were consistent with an amplified IL-23/Th17 cytokine network, a pathway long implicated in psoriasis [19]. Conversely, downregulation of epidermis development, hair-follicle cytodifferentiation, and Wnt signaling indicated loss of normal stratification and patterning, while enrichment of circadian and chromatin-related terms among downregulated genes suggested disrupted temporal and chromatin-based control. Lesional skin thus combined interferon- and cytokine-driven inflammation, antimicrobial activity, and tissue remodeling with the suppression of programs that maintain ordered differentiation and barrier integrity, consistent with current multi-omics models of psoriasis [20].

### Clinically normal patient skin occupies a transitional state

The comparison of clinically normal patient skin with healthy skin was subtler and more revealing. Although PN appears normal, its transcriptome combines basal homeostasis with early immune and stress signals, the signature of low-level subclinical activation in skin surrounding lesions. Upregulated genes were enriched for cytoplasmic ribosome and rRNA-processing terms, implying increased ribosome content and translational capacity, and for cornified envelope formation, suggesting partial engagement of late-differentiation and barrier-repair programs under inflammatory pressure. Innate immune and antimicrobial sets, including response to bacterium, leukocyte activation, and neutrophil extracellular trap formation, indicated a strengthened antimicrobial defense relative to healthy skin. Downregulated genes implied impaired lamellar lipid transport and reduced lipid synthesis, while suppression of epidermis development, hair-follicle morphogenesis, and Notch signaling pointed to weakened lineage control and incomplete epidermal maturation, reinforced by reduced cell–cell junction organization. At the regulatory level, downregulation of DNA-binding transcriptional repressors and of wound-healing regulation suggested a loosening of control over gene expression and repair. Differential expression of late cornified envelope genes further supported the role of the cornification signal. Clinically normal patient skin, therefore, occupies a transitional state, neither fully healthy nor lesional.

### Inflammation increases stepwise from healthy to lesional skin

Comparing lesions (PP) with all non-lesional skin (PN and NS combined) revealed the lesion-specific program. PP showed coordinated increases in interferon and cytokine signaling, innate antimicrobial activity, chemokine-associated recruitment, and protease regulation, together with suppression of epidermal differentiation, junctional structure, and morphogenetic control. Upregulated genes again defined an ISG-high, interferon-biased state, with canonical ISGs including *MX1*, *IFI6*, and *ISG15* among the differentially expressed genes. Concurrent enrichment of response to bacterium, chemokine receptor binding, and antimicrobial metal sequestration indicated active immune recruitment, while cornified envelope and extracellular matrix terms reflected ongoing barrier and tissue remodeling. Downregulated genes pointed to weakened epidermal layering, cohesion, and barrier assembly despite compensatory cornification signals, alongside disrupted circadian control and attenuated metabolic and hypoxia responses. Taken together with the preceding comparisons, these results describe a stepwise progression from NS through PN to PP: healthy skin shows the strongest differentiation, junctional organization, and homeostatic regulation with limited immune activation and inflammatory load, immune activation and barrier remodeling increase progressively across the three states. The data thus favor a graded continuum over a simple binary separation of healthy and diseased skin.

### The within-patient contrast isolates the lesional program

The paired comparison of PP and PN within patients confirmed interferon and cytokine signaling as the dominant lesional program, with interferon-α/β signaling, response to interferon-β, and ISG responses among the top enrichments and canonical ISGs *IFI6*, *ISG15*, and *MX1* prominently represented. Interferon signatures track disease activity across several autoimmune conditions, including psoriasis, systemic lupus erythematosus, dermatomyositis, pemphigus, and Sjögren syndrome, with type I, II, and III interferons contributing differentially to psoriatic pathophysiology [21,22]. Neutrophil degranulation and responses to bacterium and fungi indicated innate antimicrobial activity enriched for alarmins and defensins, consistent with IL-23/IL-17-driven innate immunity. Enrichment performed on the combined DEG set further highlighted responses to oxygen levels, regulation of wound healing, and cellular response to TNF, a stress- and repair-associated profile, alongside matrisome and ECM-regulator terms and vascular development programs, which are indicative of stromal remodeling and altered vasculature. Epidermal differentiation and tissue organization were represented by cornified-envelope formation, KC migration, and junction maintenance, together with regulated apoptosis and cell-cycle control, while cell activation, mononuclear cell migration, and cell–cell adhesion pointed to deregulated leukocyte trafficking at the immune–tissue interface.

### A lesion-independent disease signature

The disease-specific signature (set E), which captures genes altered in patients regardless of lesion status, exposed changes that precede visible disease. Among downregulated genes, enrichment of cornified-envelope formation and KC differentiation indicated a lesion-independent reduction in epidermal differentiation and barrier programs shared by PP and PN, and reduced antimicrobial-defense terms suggested weakened barrier defense even in clinically normal skin. The presence of leukocyte-activation terms within this downregulated component pointed to altered baseline immune regulation in skin that appears healthy. Among upregulated genes, enrichment of epidermis development, interfollicular KC differentiation, and cell-population proliferation supported a lesion-independent increase in growth- and differentiation-associated activity, while T-cell differentiation and DNA-metabolic terms indicated persistent immune priming and sustained DNA-damage and stress responses across both lesional and non-lesional areas. Enrichment of ECM organization, fibroblast migration, lipid biosynthesis, and adipocyte differentiation further implicates remodeling of the dermal microenvironment beyond the lesion itself. Complete enrichment results for all comparisons are provided in the Supplementary Materials.

### Cell-type-resolved differential expression

Cell-type-resolved differential expression in the paired set (PP vs PN) revealed a pervasive interferon and antiviral signature across cell types, underscoring the breadth of lesional inflammation. Epithelial cells were associated with epidermal development and protease regulation, and KCs separated into two functional subsets: an IL-20+ subset linked to differentiation and keratinization, and a KGF+ subset linked to proliferation, wound healing, and chemotaxis. Lesional skin thus combined a pervasive interferon environment with active keratinocyte remodeling, consistent with established psoriasis pathology [23]. Cell type assignments are shown in Fig 3, with additional panels in the Supplementary Materials.

**Fig 3.**
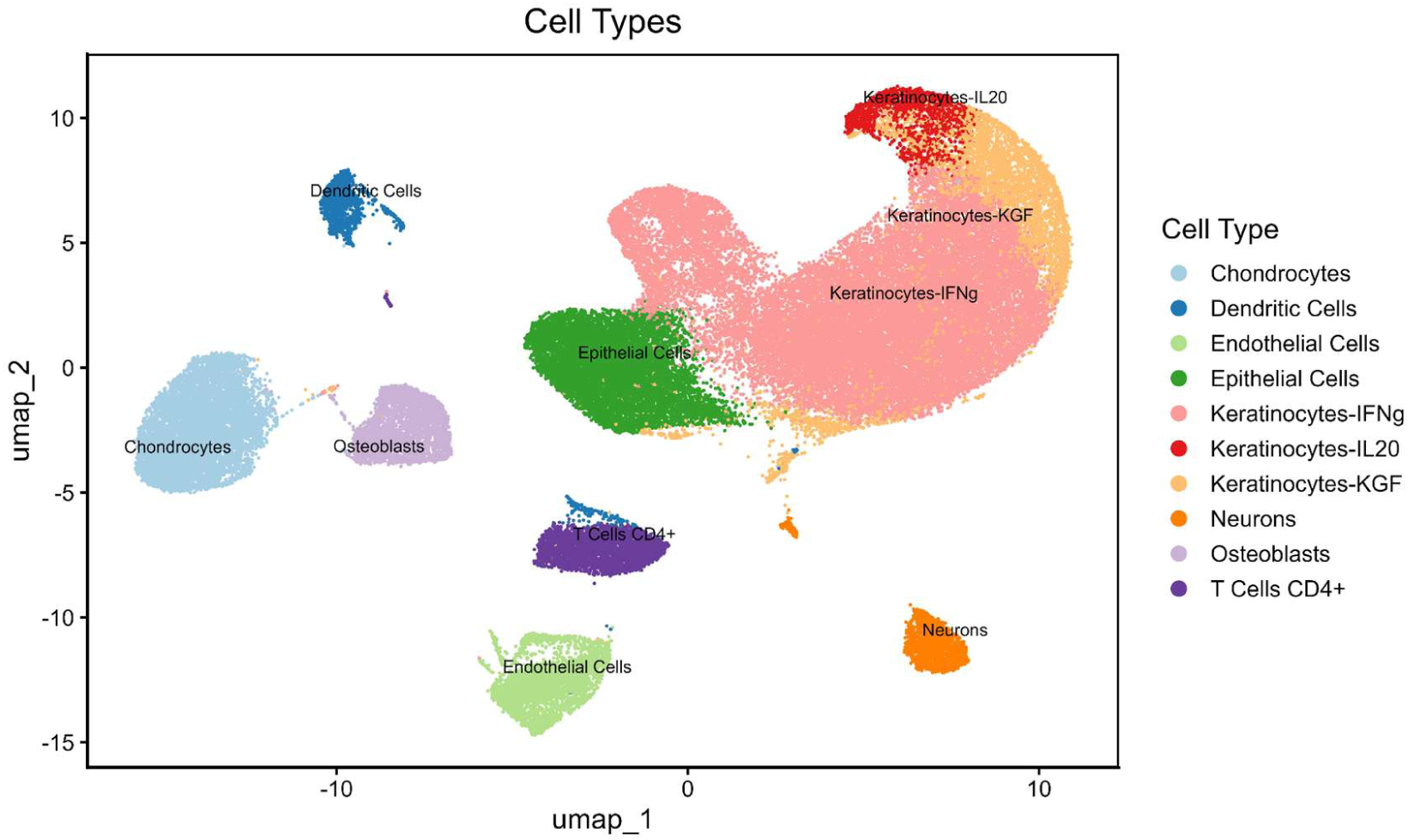
Cell-type assignments for the paired PN–PP set. Cells are shown in UMAP space after Harmony integration and colored by the cell-type label assigned with SingleR and checked against established literature markers.

### Pseudotime resolves an ordered continuum of transcriptional states

To move from static contrasts to dynamics, we ordered cells along pseudotime. Pseudotime analysis arranges individual cells along a continuous axis that reflects a biological process such as differentiation or activation, inferred from transcriptional state rather than elapsed time, with higher values indicating more advanced states within a cell type [18]. The number of genes correlated with pseudotime varied by cell type, with 1,020 in endothelial cells, 509 in IL-20+ KCs, and 142 in KGF+ keratinocytes (Table 1), indicating that a defined gene subset dominates cell ordering in each lineage; full lists of positively and negatively correlated genes are provided in the Supplementary Materials. For each cell type, pseudotime distributions were examined, and, where trajectories branched, branches containing cells from multiple tissues were prioritized so that transcriptional transitions between paired-set samples could be observed directly. Node-anchored differential expression then contrasted the most divergent populations along each branch, excluding intermediate cells.

**Table 1.** Number of genes correlated with pseudotime. Genes were retained at an absolute Spearman correlation coefficient above 0.1 and an adjusted p-value below 0.01.

| Cell Type | Positively Correlated Genes | Negatively Correlated Genes | Total Correlated Genes |
| --- | --- | --- | --- |
| Endothelial cells | 429 | 591 | 1,020 |
| KCs: IL20 | 370 | 139 | 509 |
| KCs: KGF | 94 | 48 | 142 |

### Endothelial cells mature from an inflamed toward a stabilized state

Endothelial cells traced a trajectory from a patient-normal origin toward lesional states (Fig 4). The starting region, E1, was composed predominantly of PN cells. Relative to E2 and E3, E1 showed lower expression of endothelial differentiation and blood-vessel development programs and higher expression of programs supporting balanced vascularization, immune regulation, stress response, and cell turnover, functions typically lost in lesions, where their dysregulation is reflected in defective angiogenesis, persistent inflammation, and KC hyperproliferation; focal-adhesion processes were also lower in E1. The E2 and E3 regions, by contrast, showed higher expression of genes associated with hyper-inflammatory, hyperproliferative, and pro-angiogenic states, resembling a non-healing wound with heightened immune and vascular activity, together with lower expression of protective, regulatory, and apoptotic pathways. The genes correlated with pseudotime described a different axis. Adhesion-molecule and MHC-II binding, leukocyte adhesion, cytokine response, VEGFR2 signaling, the RHO-GTPase cycle, chromatin remodeling, ECM organization, motility, angiogenesis, and blood-vessel development were all negatively correlated with pseudotime, whereas metabolic and translational programs were positively correlated, so that increasing pseudotime tracked a shift from an inflamed, adhesive, angiogenic profile toward a more metabolically active, barrier-associated one. This pseudotime axis is therefore not equivalent to the PN-versus-PP contrast between E1 and E2/E3, and the two should not be read as a single disease-progression axis. We therefore interpret these results conservatively: endothelial cells occupied a continuum of inflammatory, angiogenic, and metabolically mature states, but the inferred trajectory was not strictly aligned with a unidirectional PN-to-PP transition. Distinguishing whether these represent separate branches or a single ordering would require an explicitly justified root and independent validation.

**Fig 4.**
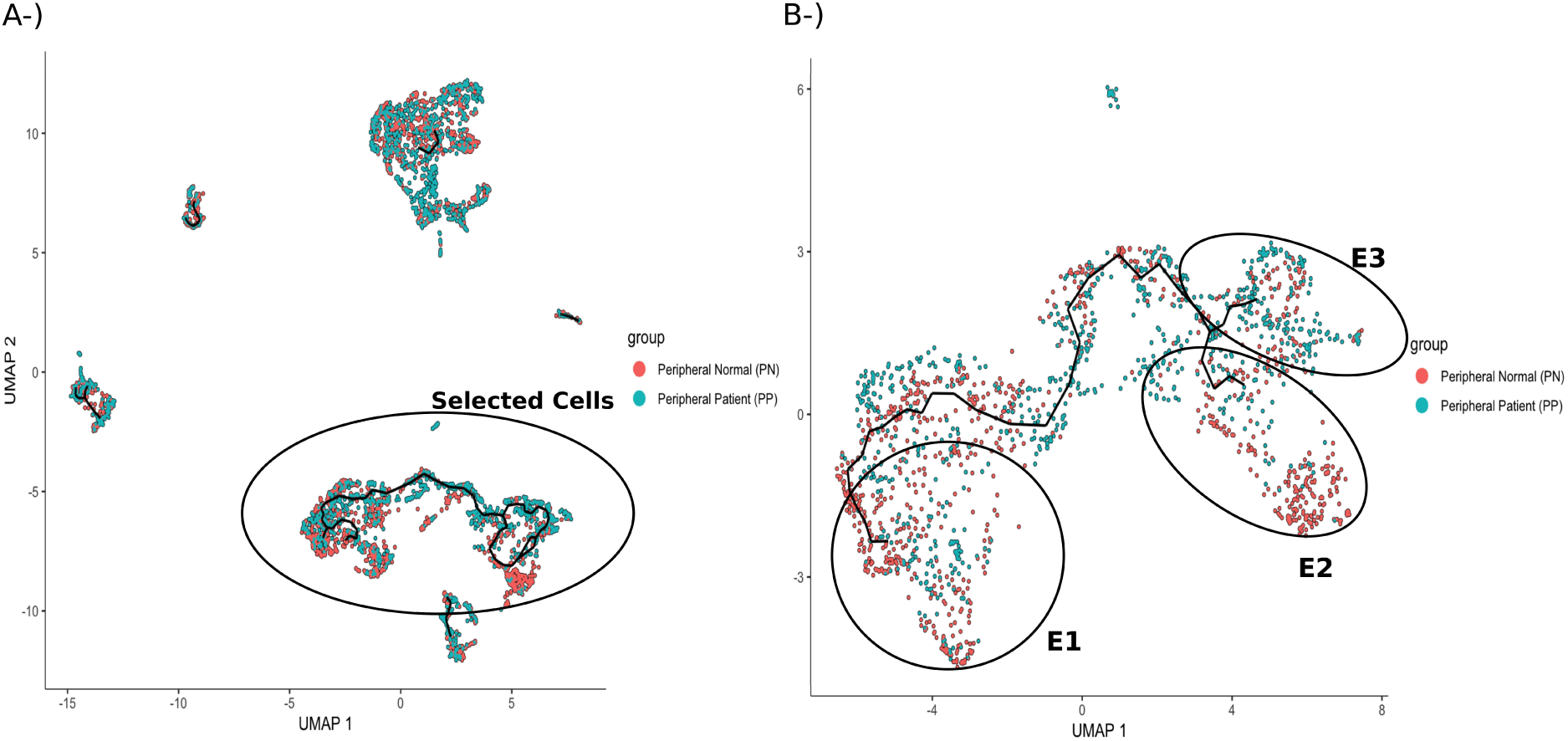
Inferred trajectory for endothelial cells in the paired set. (A) Cells are shown in UMAP space with the Monocle3 principal graph overlaid in black. (B) Circled areas indicate the node-anchored regions selected for differential expression (E1, E2, E3); intermediate cells were excluded from these comparisons. PN, non-lesional psoriatic skin; PP, lesional psoriatic skin.

### IL-20+ keratinocytes shift from protective toward hyperproliferative states

IL-20+ KCs resolved into three regions (Fig 5): IL1, dominated by PN cells; IL3, dominated by PP cells; and a central region (IL2) of mixed origin. The IL1 region, closest to normal tissue, comprised metabolically active cells engaged in protein synthesis and barrier support, with intact stress-response and apoptosis regulation, in contrast to the hyperproliferative lesional cells; these cells participated in anti-inflammatory and homeostatic pathways that may buffer against full psoriatic transformation, although vascular endothelial growth factor (VEGF) activity indicated that skin adjacent to lesions remains susceptible to angiogenic and inflammatory change. In the central, mixed region, KCs maintained barrier components, including cornified-envelope formation, desmosomes, and lamellar bodies, while adapting to stress and hypoxia through increased *HMOX1* and oxygen-response expression, raising energy and ion-transport activity, evading natural killer cell cytotoxicity, and remodeling lipid and endoplasmic reticulum stress responses, consistent with impaired but ongoing differentiation; this region thus balanced barrier maintenance against stress adaptation under inflammatory and metabolic pressure. The IL3 region, dominated by lesional tissue, upregulated programs of hyperproliferation, protein synthesis, metabolic reprogramming, mitochondrial stress, dysfunctional barrier formation, excessive antimicrobial and neutrophil activation, and chronic stress relative to IL1 and IL2. Along pseudotime, IL-20+ KCs, therefore, moved from an early, inflamed, stress-engaged state toward a metabolically rewired, proteostatic one: genes negatively correlated with pseudotime implicated antimicrobial responses, vitamin D receptor signaling, and KC differentiation, whereas positively correlated genes emphasized oxidative phosphorylation, translation and ribosome biogenesis, chaperone-mediated trafficking, autophagy, and lipid and carbon catabolism, consistent with cytokine activation followed by cellular recovery and stabilization. Collectively, lesional IL-20+ KCs define a hyperproliferative, metabolically overactive, immune-stimulated state that drives plaque formation and maintenance.

**Fig 5.**
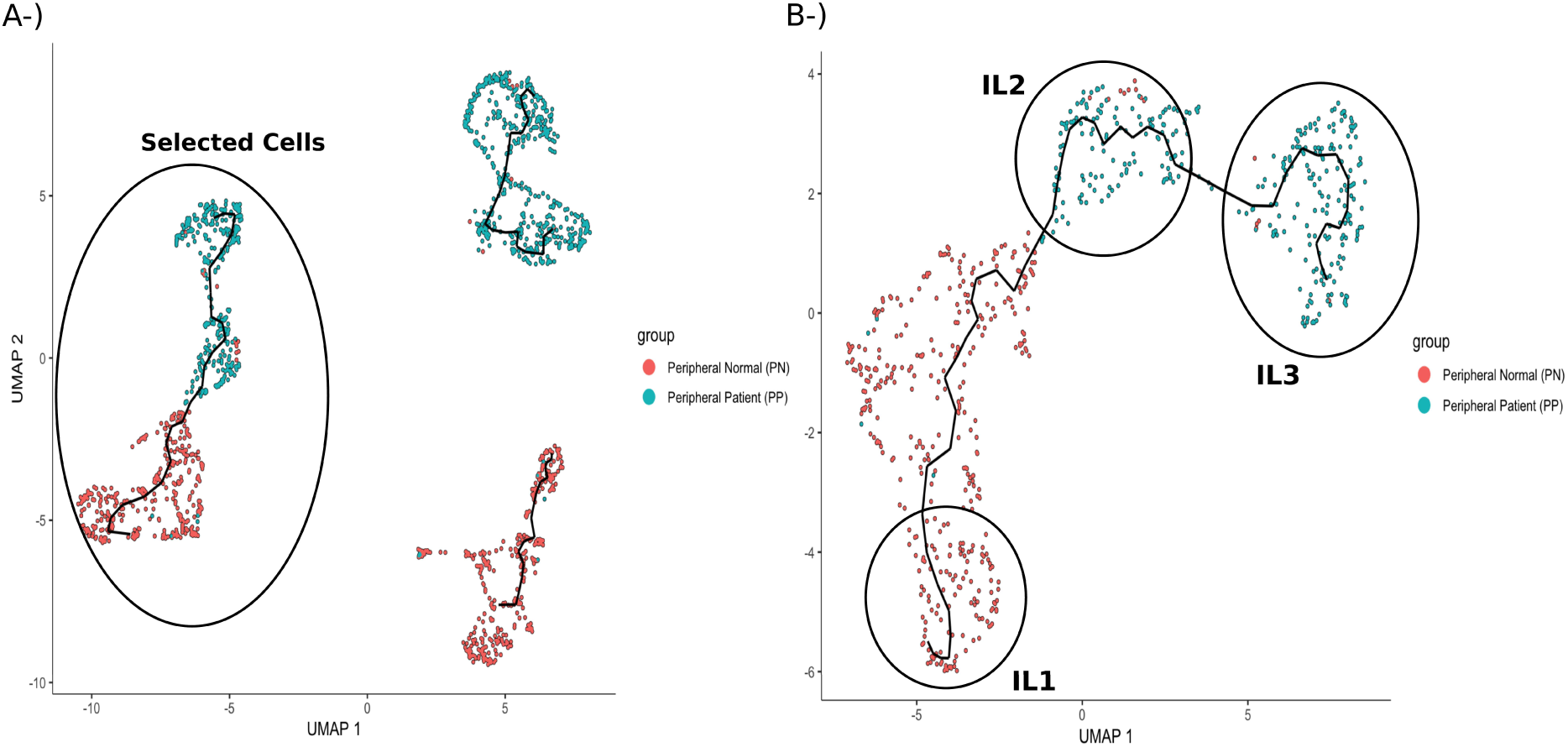
Inferred trajectory for KCs: IL20 cells in the paired set. (A) Cells are shown in UMAP space with the Monocle3 principal graph overlaid in black. (B) Circled areas indicate the node-anchored regions selected for differential expression (IL1, IL2, IL3); intermediate cells were excluded from these comparisons. PN, non-lesional psoriatic skin; PP, lesional psoriatic skin.

### KGF+ keratinocytes converge on a common lesional phenotype

KGF+ KCs showed a distinct distribution, with a trajectory extending from healthy-associated tissue into two branches composed mainly of lesional cells (Fig 6). The K1 region, enriched for healthy-tissue cells, upregulated ribosomal activity and downregulated differentiation, chromatin regulation, and adhesion programs relative to K2 and K3, suggesting a latent, pre-activated state primed for psoriatic transformation under inflammatory conditions. The K2 and K3 regions each upregulated ribosomal and metabolic machinery supporting hyperproliferation while downregulating differentiation, adhesion, and structural programs, pointing to barrier dysfunction and epidermal disorganization; loss of transcriptional and hormonal regulation implied uncontrolled growth and diminished anti-inflammatory capacity, and upregulation of genes linked to mitochondrial imbalance reflected unmet energy demands and oxidative stress. Lesional KGF+ KCs were therefore highly active in growth and protein synthesis yet deficient in differentiation, cell-cycle control, and stress response, a combination that favors thick, inflamed, poorly differentiated plaques. Notably, although K2 and K3 lie on different branches, they share similar DEGs, indicating convergence of KGF+ KCs onto a common psoriatic phenotype from distinct trajectory positions and a robust pathological program that persists across spatial and temporal contexts. Pseudotime analysis further showed a shift toward terminal differentiation and barrier strengthening, with upregulation of cornified-envelope formation, KC differentiation, antimicrobial metal sequestration, cytoskeletal remodeling, mitochondrial respiration, proteasome activity, and epidermal-barrier development, while genes negatively correlated with pseudotime indicated that early translational-defense and cytoprotective programs diminish in cells committed to cornification and stable barrier function (Supplementary Materials).

**Fig 6.**
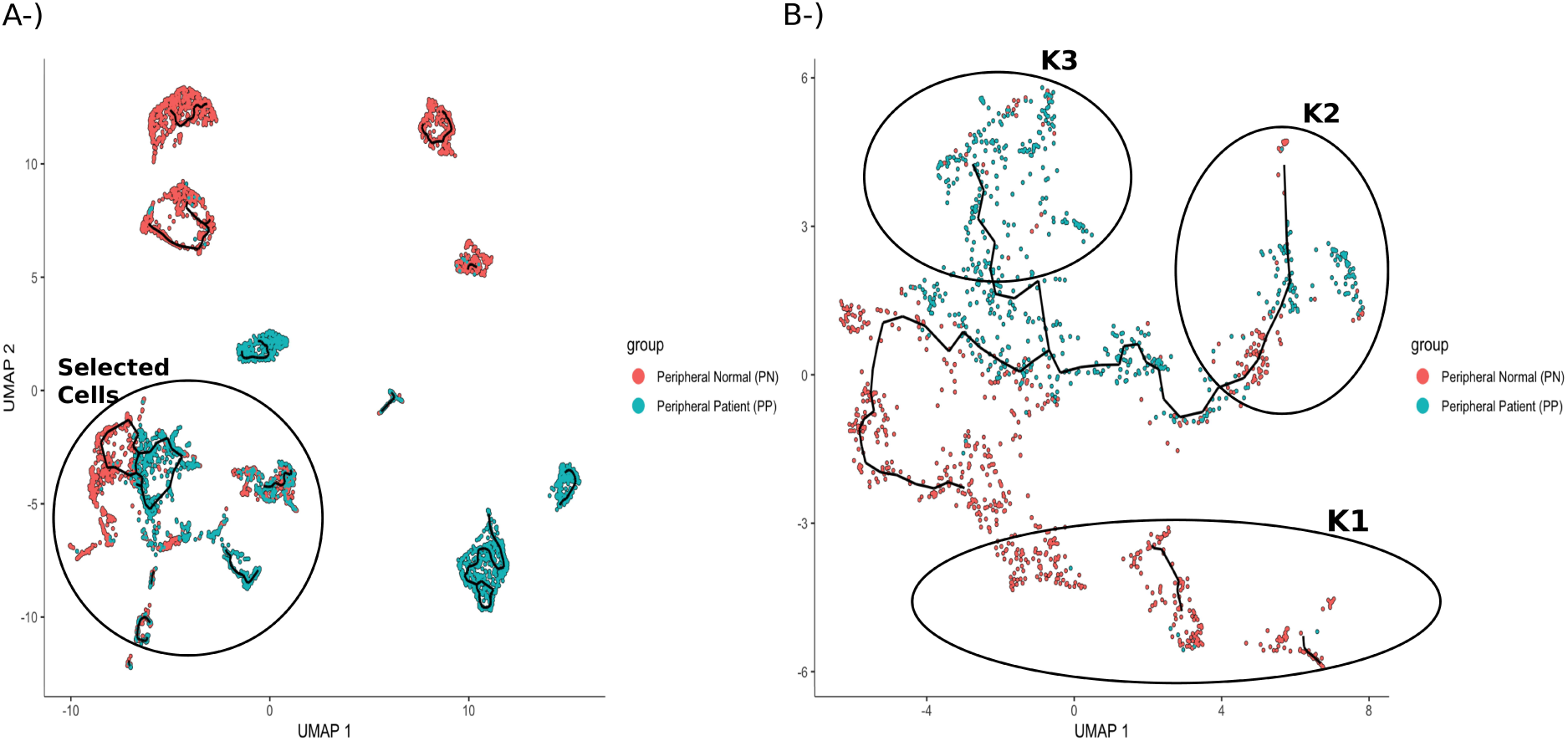
Inferred trajectory for KCs: KGF cells in the paired set. (A) Cells are shown in UMAP space with the Monocle3 principal graph overlaid in black. (B) Circled areas indicate the node-anchored regions selected for differential expression (K1, K2, K3); intermediate cells were excluded from these comparisons. PN, non-lesional psoriatic skin; PP, lesional psoriatic skin.

### Neurons show no coordinated transition

Neurons also displayed a pseudotime trajectory with three regions, from N1 to N3. The proportion of PP cells increases from the PN-dominated N1 region toward the more evenly mixed N3 (Fig 7). Unlike the other cell types, however, differential expression analysis between branches yielded no enriched biological processes, indicating an absence of coordinated transcriptional differences between PP and PN within the neuronal compartment.

**Fig 7.**
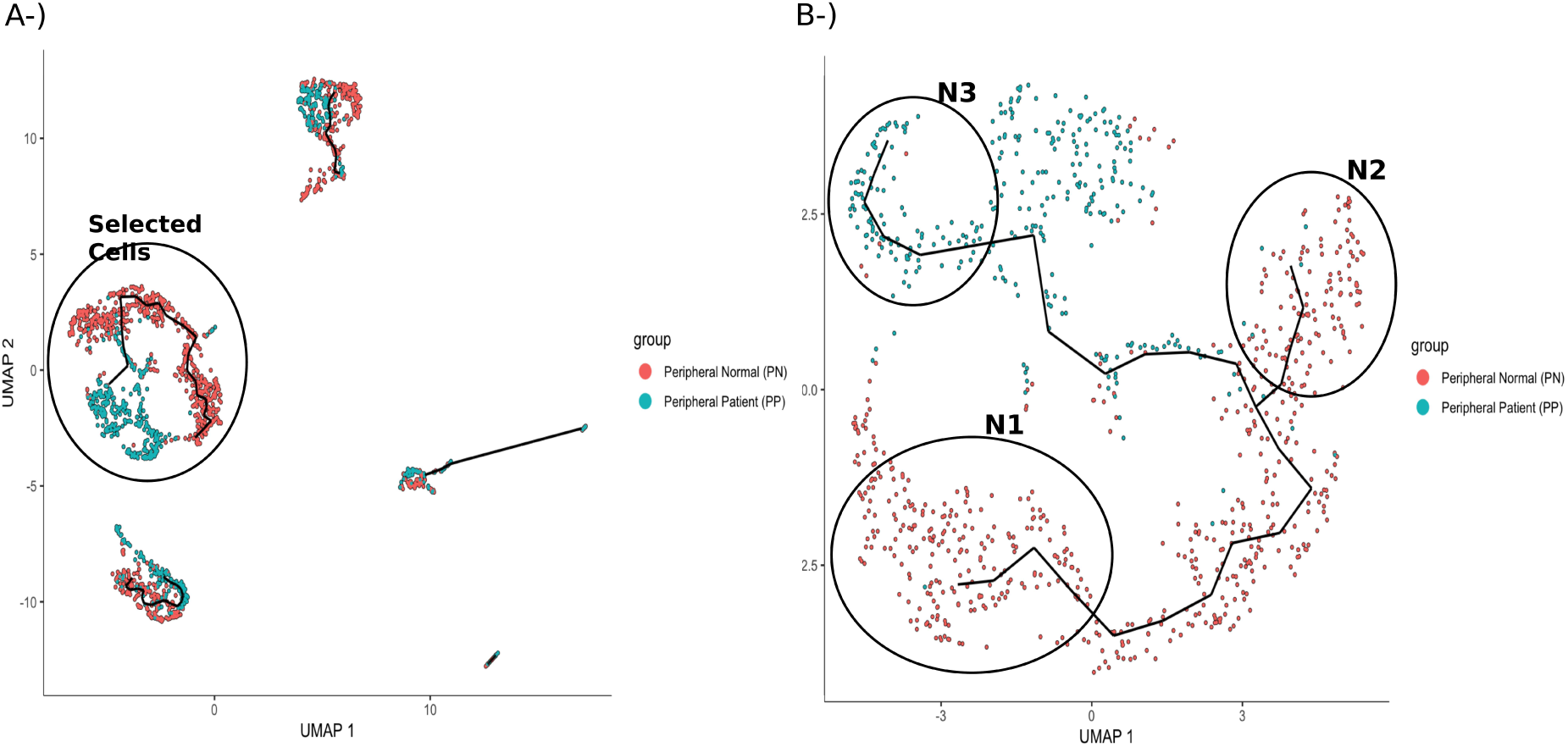
Inferred trajectory for neuron cells in the paired set. (A) Cells are shown in UMAP space with the Monocle3 principal graph overlaid in black. (B) Circled areas indicate the node-anchored regions compared (N1, N2, N3); intermediate cells were excluded from these comparisons. PN, non-lesional psoriatic skin; PP, lesional psoriatic skin.

Consequently, this negative result serves as a critical baseline control, confirming that our node-anchored trajectory model selectively identifies authentic pathological transitions in responding structural cell populations such as endothelial cells and keratinocytes, rather than forcing artificial transcriptional gradients across all captured cell types.

### A coordinated endothelial–keratinocyte continuum

Across cell types, these trajectories describe an ordered continuum of transcriptional states associated with clinically normal and lesional skin from the same patients. Beyond cataloguing disease states, this framework provides a basis for investigating regional heterogeneity and why lesions occur in some areas while neighboring skin remains clinically normal, a question that will require spatial and longitudinal data to address directly.

## Conclusions

Differential expression across single-cell profiles of psoriatic and healthy skin identified genes consistently altered in patients regardless of lesion status. The disease-specific signature revealed lesion-independent disruption of the epidermal barrier and innate immunity, with downregulation of cornified-envelope formation, KC differentiation, and antimicrobial defense, and with leukocyte-activation terms pointing to altered baseline immune regulation in clinically normal skin. Persistent immune priming and cellular stress were reflected in upregulated epidermal growth, keratinocyte proliferation, T-cell differentiation, stress response, and DNA-metabolic programs, while ECM organization, fibroblast migration, and lipid biosynthesis indicated dermal and metabolic remodeling beyond the lesion.

Integrating these contrasts with pseudotime and branch-specific analyses yields a coherent, cell-type-resolved model of progression in which endothelial cells and keratinocyte subsets undergo structured, trajectory-dependent transitions. Endothelial cells advance from an early, inflamed, adhesive state toward a metabolically mature, barrier-stabilized phenotype, reflecting a shift from active vascular remodeling toward partial resolution within the lesion. In parallel, IL-20+ and KGF+ KCs undergo strong pseudotime-dependent reprogramming, moving from pre-activated or homeostatic stages in clinically normal skin toward hyperproliferative, metabolically rewired, stress-adapted states in lesions. These keratinocyte paths converge on a pathogenic program of epidermal thickening, barrier dysfunction, and plaque maintenance, characterized by excessive protein synthesis, mitochondrial and endoplasmic reticulum stress, weak differentiation, and persistent inflammatory signaling. Intermediate cells, present in both lesional and clinically normal skin, retain homeostatic and protective features and may delay full disease progression, marking points of plasticity in the trajectory. The behavior of pseudotime-correlated genes supports each of these transitions.

This continuum-based view frames psoriasis as a multicellular process involving coordinated endothelial and keratinocyte transcriptional change, and is consistent with the observation that clinically normal patient skin already carries a partial disease-associated signature. Rather than demonstrating a literal temporal sequence, our analysis identifies an ordered continuum of transcriptional states across healthy, non-lesional, and lesional skin, with distinct disease-associated trajectories in endothelial and keratinocyte populations. The intermediate states identified here are candidates for longitudinal or spatial validation and, if confirmed, for investigation as points at which disease-associated change might be interrupted.

Several limitations should be noted. First, the data are cross-sectional: pseudotime orders cells by transcriptional similarity and cannot establish that individual non-lesional cells develop into lesional cells, so all trajectory interpretations describe ordered transcriptional states rather than observed temporal sequences. Second, the study is based on a single publicly available dataset of matched lesional and clinically normal skin together with healthy controls; additional patients, samples, and independent datasets would strengthen reproducibility, enable a fuller assessment of inter-patient heterogeneity, and improve statistical power. Third, cell-type annotation relies on reference-based prediction together with literature markers, so subset labels, including the IL-20-high and KGF-high keratinocyte designations, carry annotation uncertainty and would benefit from explicit marker-level validation. Fourth, trajectory results depend on the choice of root and on which branches are analyzed; alternative roots or branch selections could yield different orderings, and prioritizing branches containing cells from more than one tissue may introduce selection bias. Fifth, differential expression was computed at the level of individual cells, which does not treat the sample or patient as the unit of biological replication; sample-aware or pseudobulk models would provide more conservative inference, particularly for the paired PP-PN contrast. Finally, the findings are transcriptional and unvalidated: no spatial, protein-level, or functional experiments were performed to confirm the inferred states, their tissue localization, or their role in lesion development.

## Data Availability

The dataset used in this study can be downloaded from the NCBI GEO repository under accession number GSE173706.

## Author Contributions

G.U.: methodology, study design, investigation, data curation, writing—original draft, and visualization. D.K.: methodology, study design, investigation, data curation, writing—original draft, and visualization. H.K.: conceptualization, study design, supervision, writing—review and editing. P.P.: conceptualization, study design, supervision, writing—review and editing.

## Statements and Declarations

This study is a secondary analysis of previously published, de-identified single-cell RNA-seq data (GSE173706); no new human participants’ approval was required for this work. Ethical approval and participant consent for the original sample collection are described in the primary studies [4,8].

