## Supplementary Figures for "A transcriptional continuum from clinically normal to lesional skin: Single-cell trajectory analysis of psoriasis"

**Metascape Enrichment Results**

**NS = Healthy Normal Skin**

**PP = Lesional Psoriatic Skin**

**PN = Peripheral Normal Skin from Psoriasis Patients**

**Paired and Complete Enrichment Results**

Dimension reduction was performed using Principal Component Analysis (PCA) with two different parameter settings. For the complete dataset, the first 15 principal components (PCs) were used for clustering and downstream differential expression gene (DEG) analysis, whereas for the paired dataset, analyses were conducted using the first 10 PCs. The complete dataset contained a larger number of samples and cells, resulting in greater overall variability and therefore requiring more PCs to capture a comparable proportion of the total variance relative to the paired subset.

**The Complete Set**

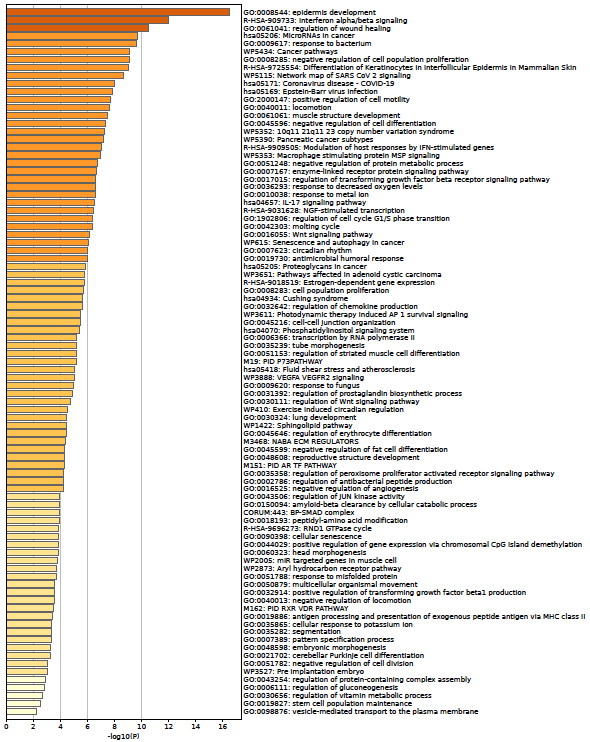

**Fig 1**. Enrichment results for NS vs all.

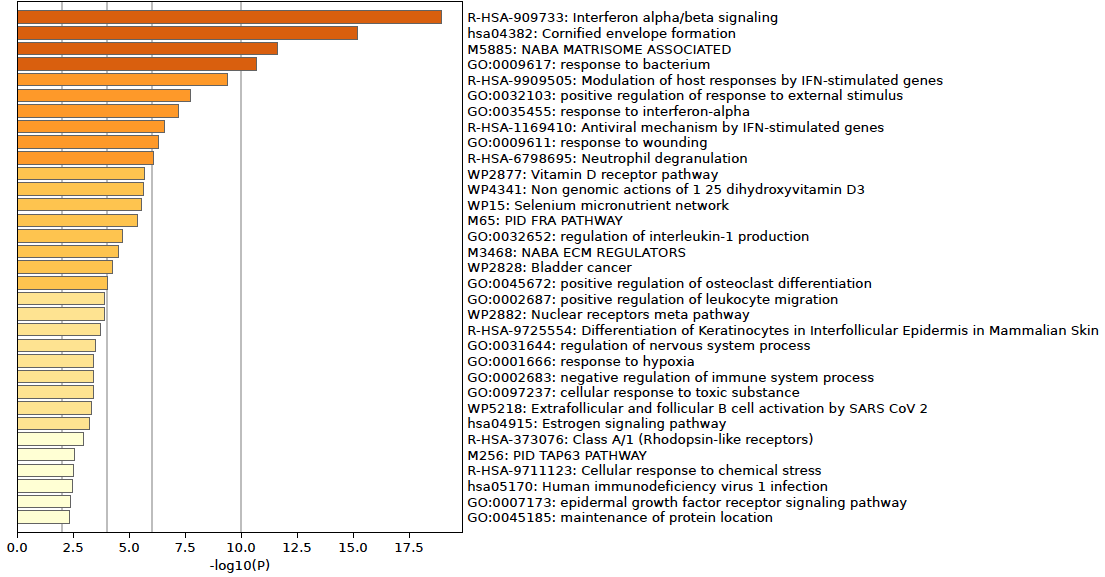

**Fig 2.** Enrichment results for PN vs all.

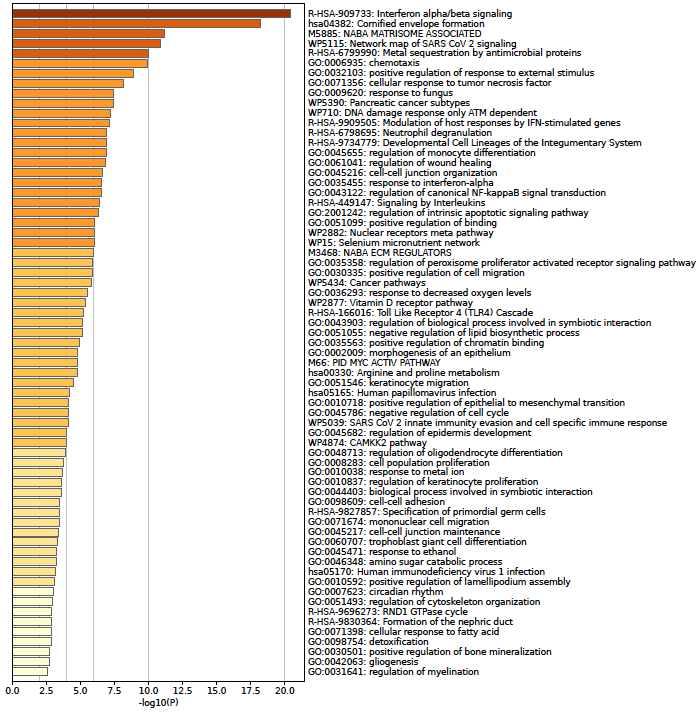

**Fig 3.** Enrichment results for PP vs all.

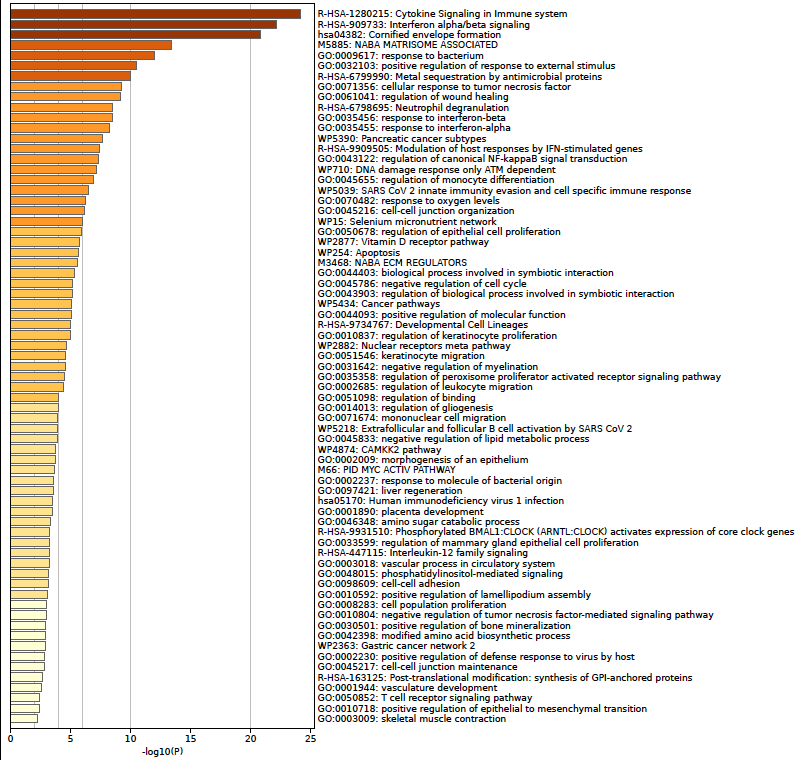

**Fig 4.** Enrichment results for PP vs PN.

**The Paired Set**

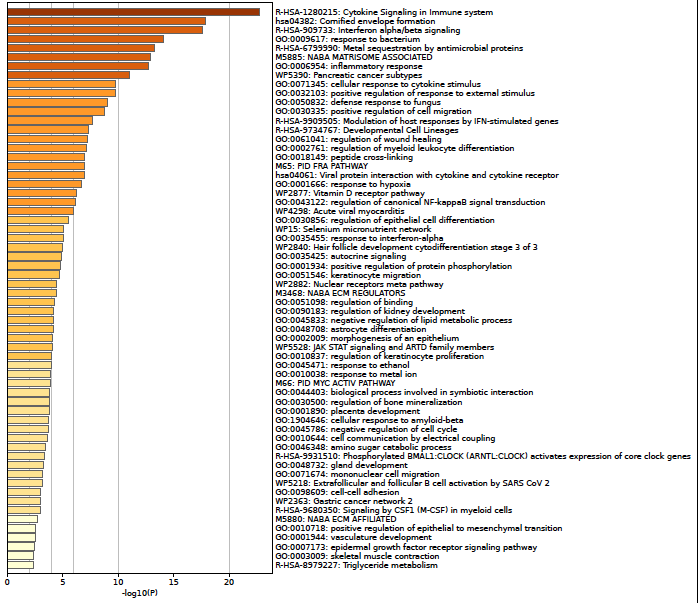

**Fig 5.** Enrichment results for PP vs PN.

**Enrichment Results for both Upregulated and Downregulated Genes for Complete Set**

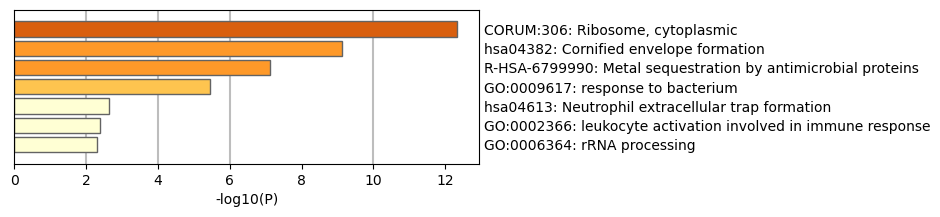

**Fig 6.** Upregulated genes enrichment results for PN vs NS.

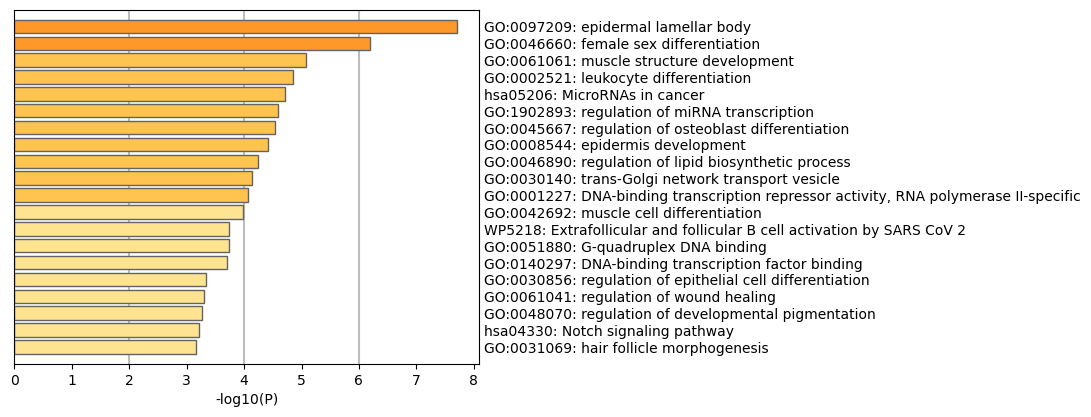

**Fig 7.** Downregulated genes enrichment results for PN vs NS.

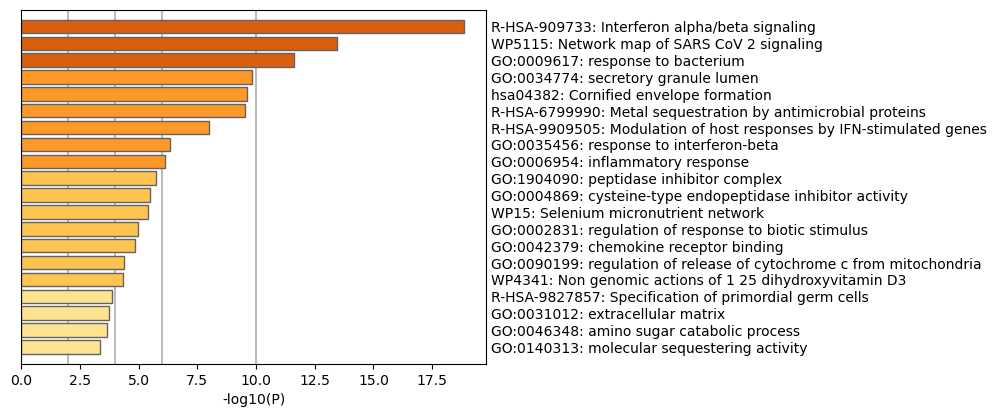

**Fig 8.** Upregulated genes enrichment results for PP vs all.

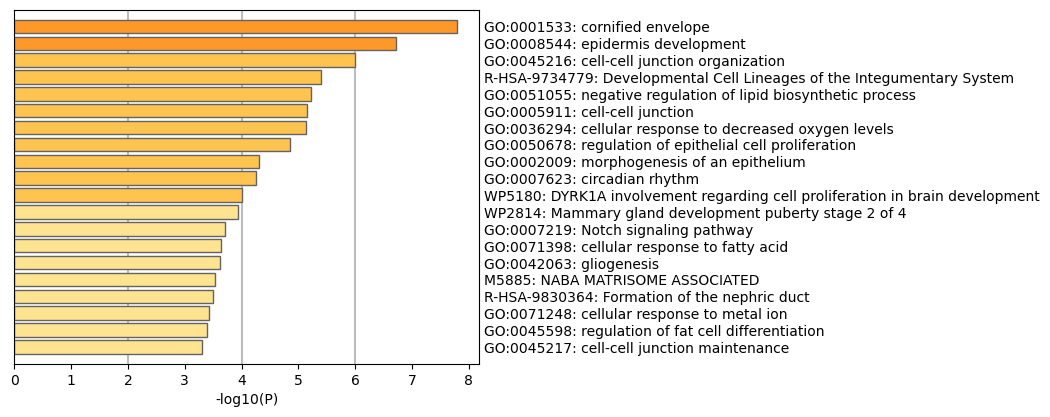

**Fig 9.** Downregulated genes enrichment results for PP vs all.

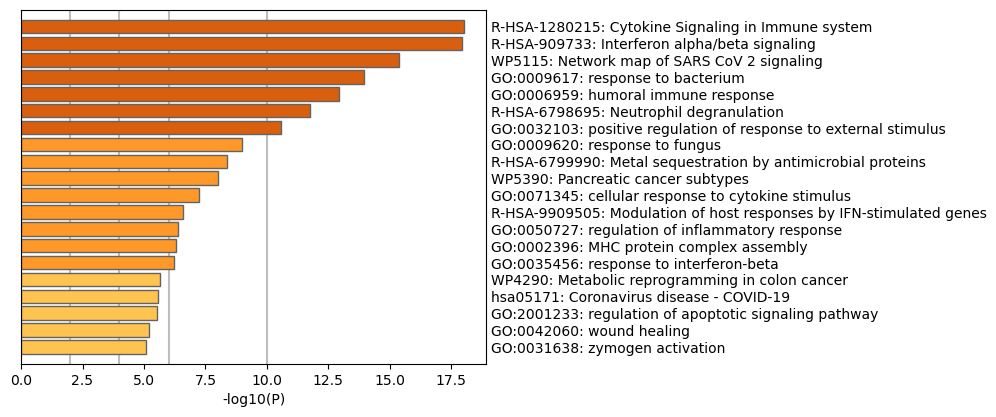

**Fig 10.** Upregulated genes enrichment results for PP vs NS.

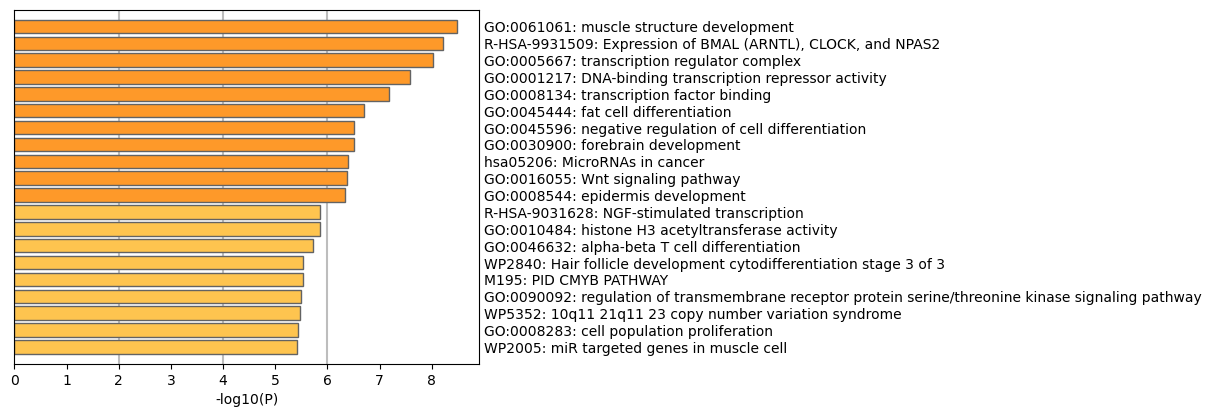

**Fig 11.** Downregulated genes enrichment results for PP vs NS.

**Positively and Negatively Correlated Genes on Pseudotime Enrichment Results**

**Negatively Correlated**

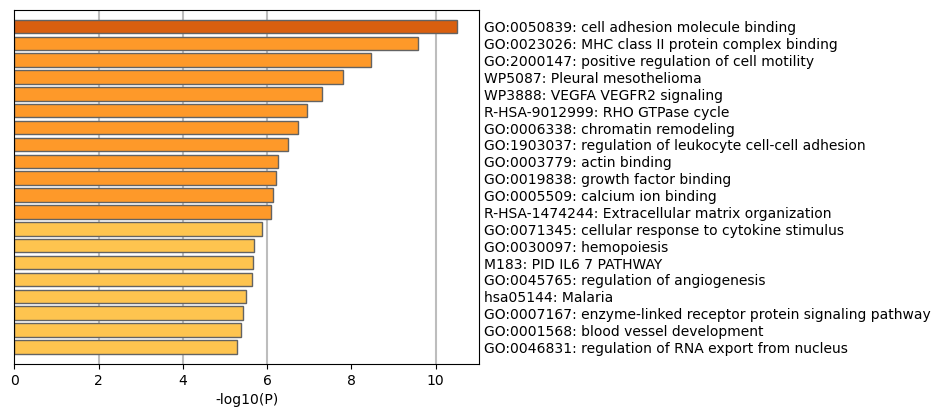

**Fig 12.** Negatively correlated genes enrichment results for endothelial cells.

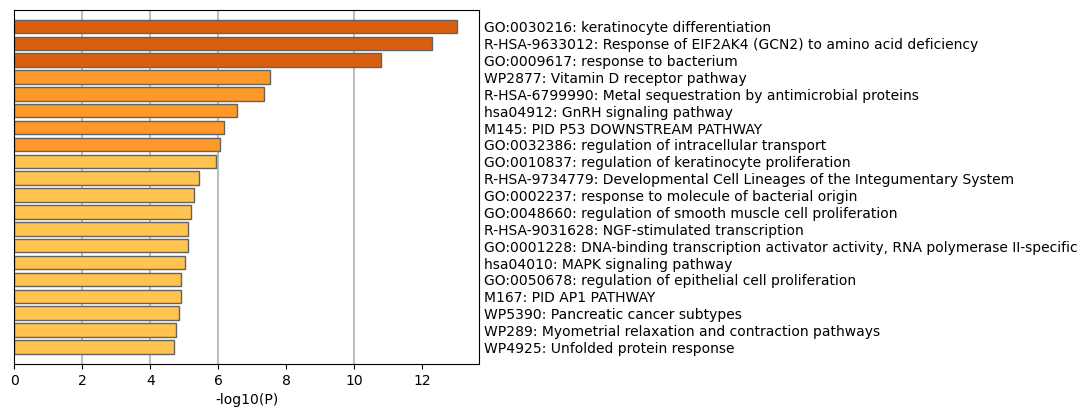

**Fig 13.** Negatively correlated genes enrichment results for KCs: IL20 cells.

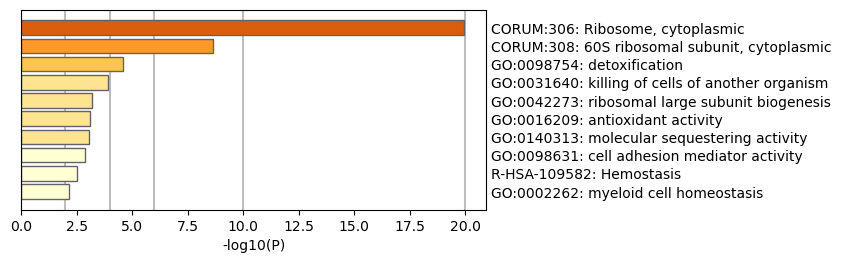

**Fig 14.** Negatively correlated genes enrichment results for KCs: KGF cells.

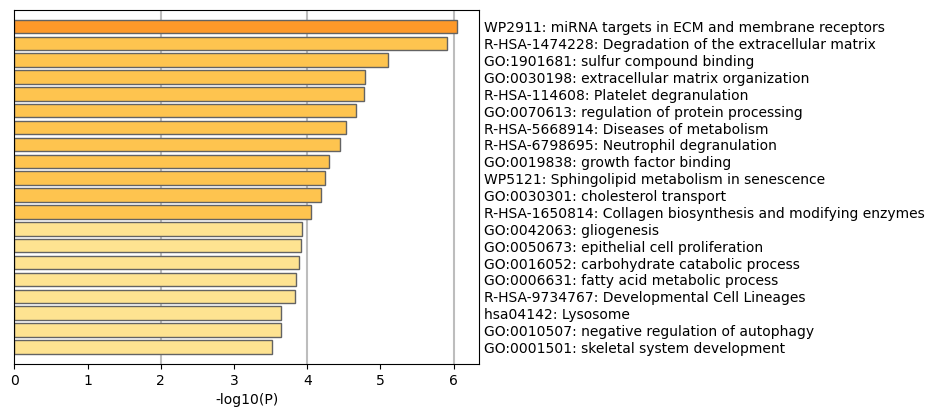

**Fig 15.** Negatively correlated genes enrichment results for neuron cells.

**Positively Correlated**

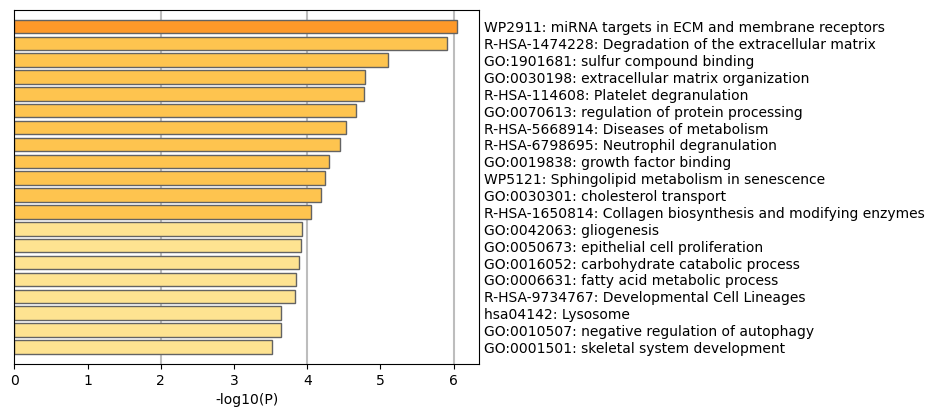

**Fig 16.** Negatively correlated genes enrichment results for endothelial cells.

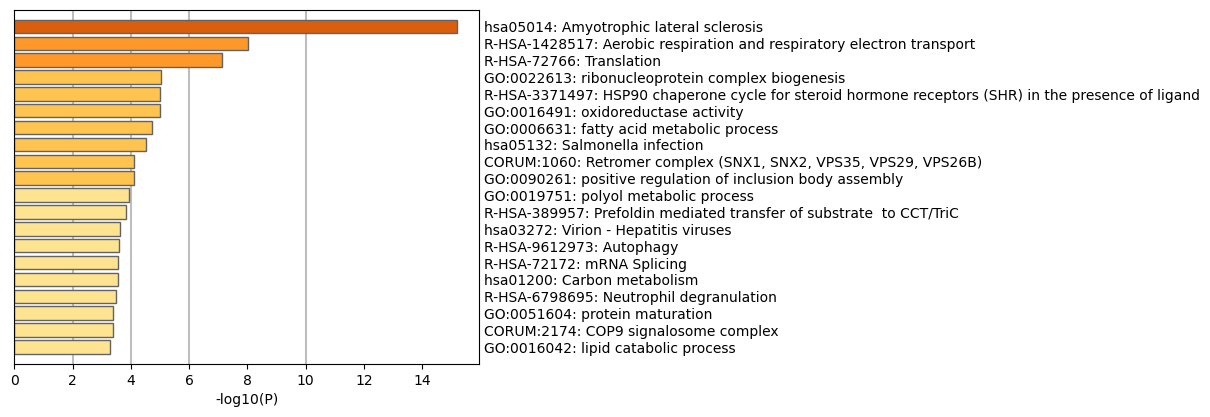

**Fig 17.** Positively correlated genes enrichment results for KCs: IL20 cells.

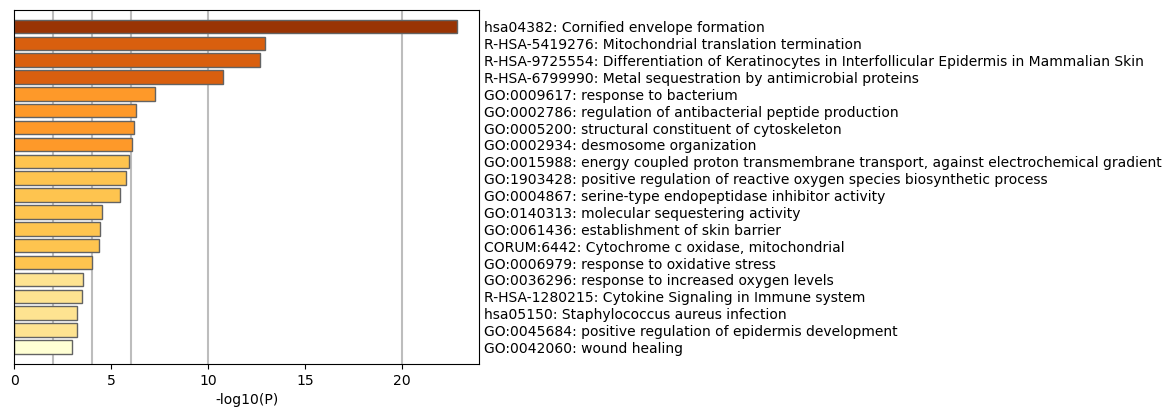

**Fig 18.** Positively correlated genes enrichment results for KCs: KGF cells.

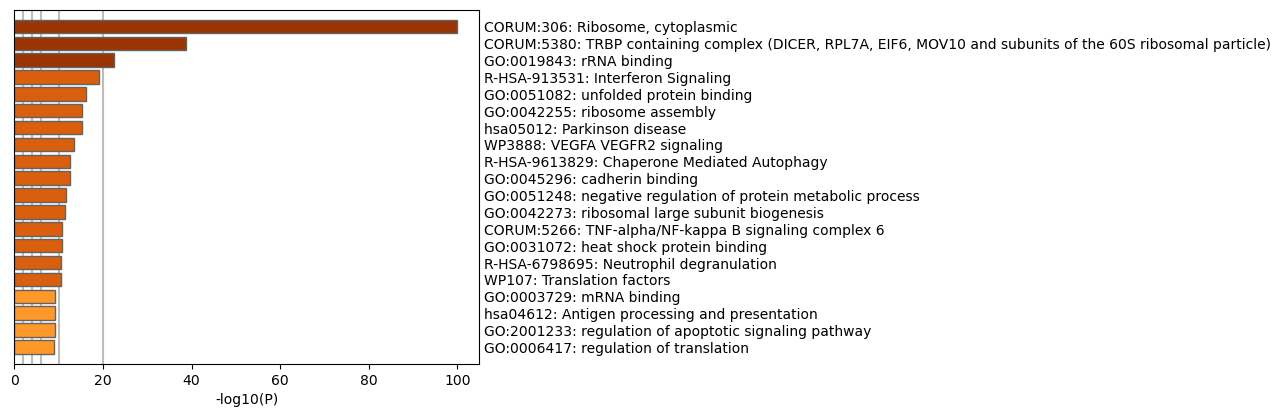

**Fig 19.** Positively correlated genes enrichment results for neuron cells.

**Pseudotime enrichment results based on cell type**

**Endothelial Cells**

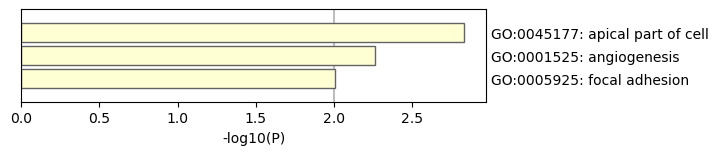

**Fig 20.** Downregulated genes enrichment results for E1 region.

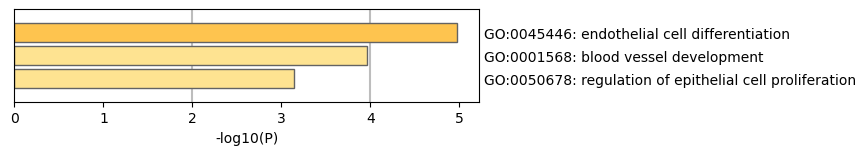

**Fig 21.** Downregulated genes enrichment results for E2 region.

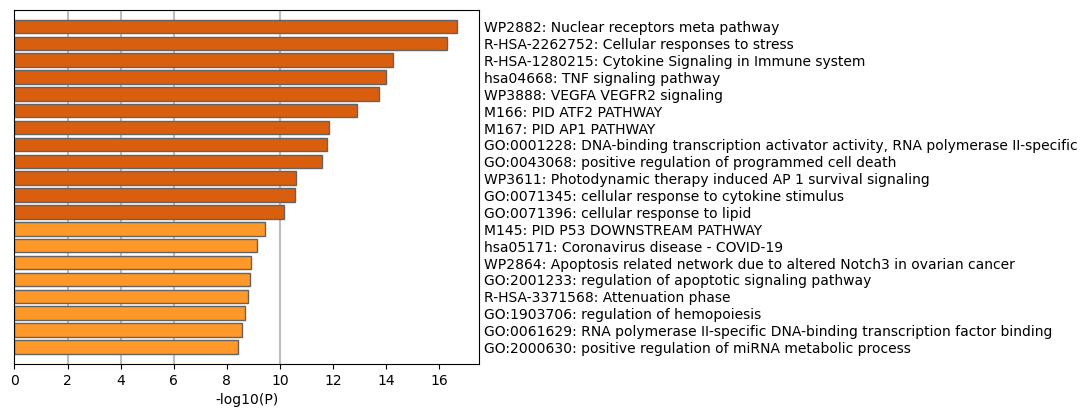

**Fig 22.** Downregulated genes enrichment results for E3 region.

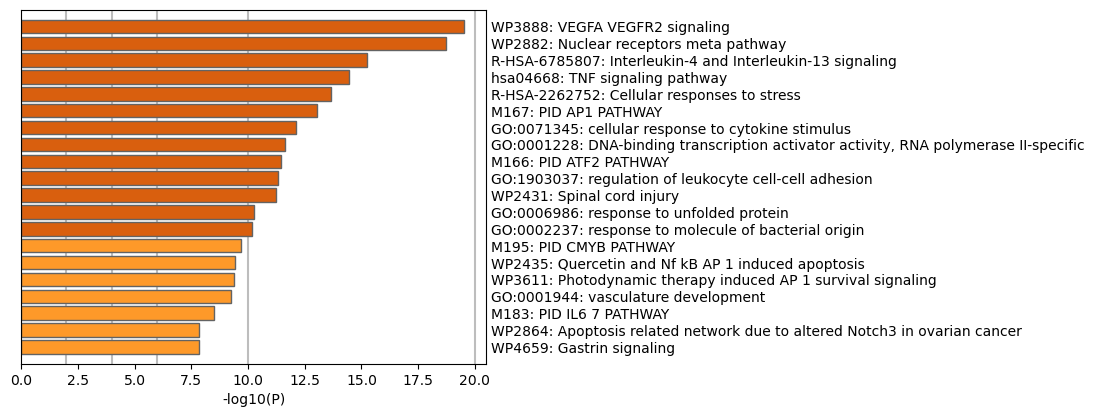

**Fig 23.** Upregulated genes enrichment results for E1 region.

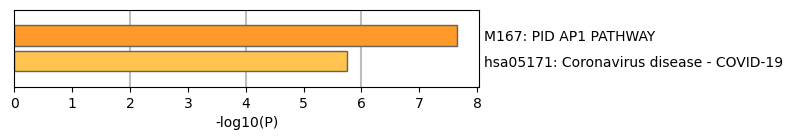

**Fig 24.** Upregulated genes enrichment results for E2 region

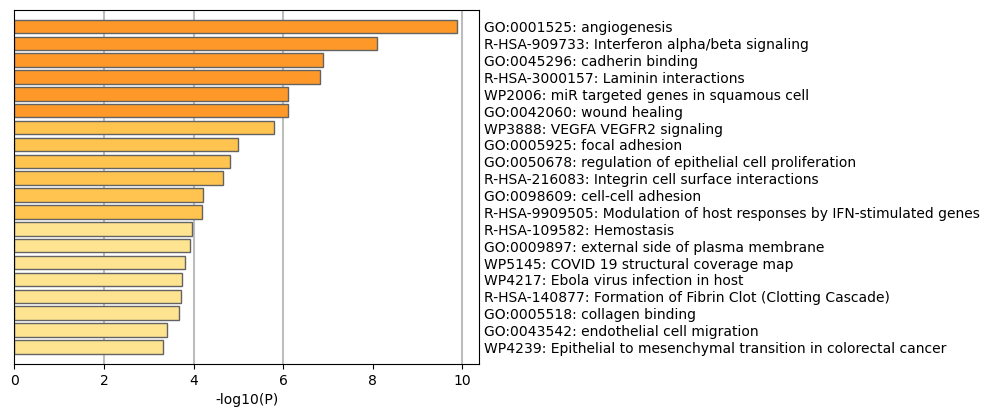

**Fig 25.** Upregulated genes enrichment results for E3 region.

**KCs: IL20 Cells**

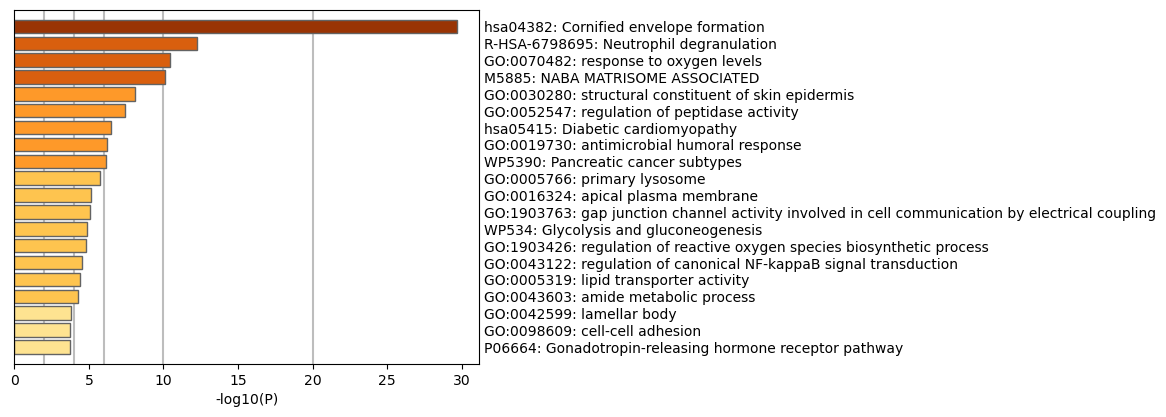

**Fig 26.** Downregulated genes enrichment results for IL1 region.

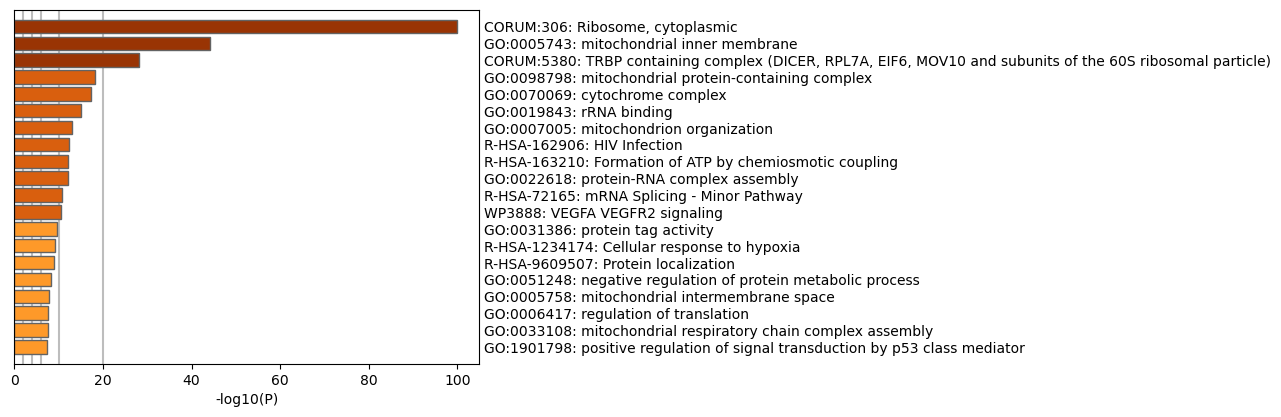

**Fig 27.** Downregulated genes enrichment results for IL2 region.

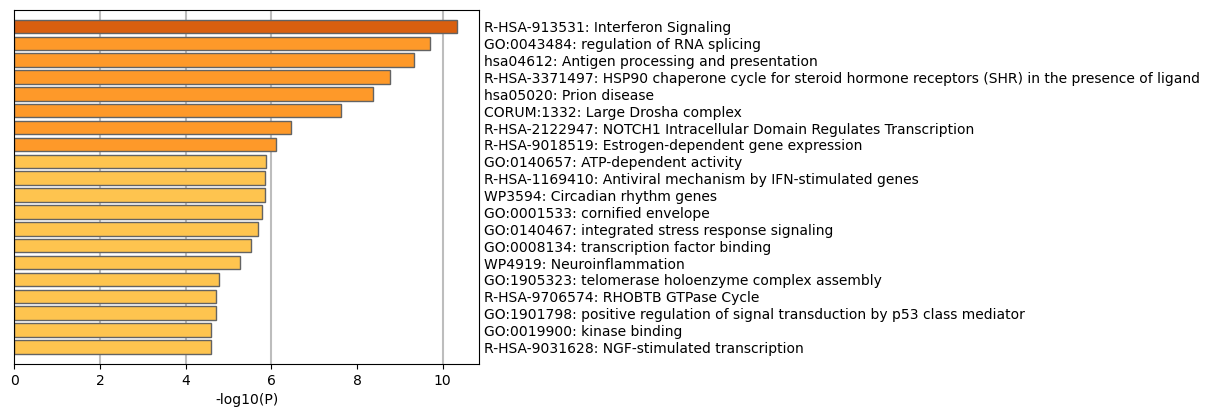

**Fig 28.** Downregulated genes enrichment results for IL3 region.

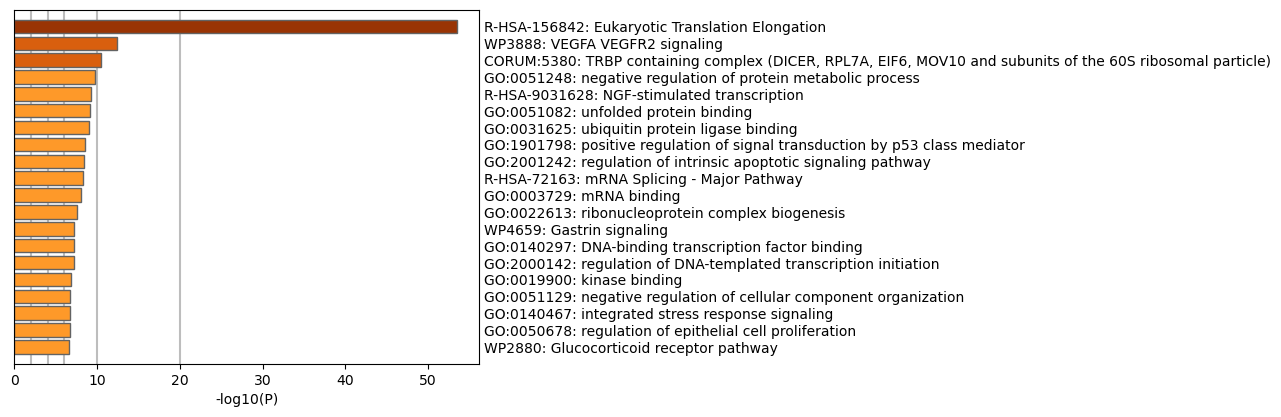

**Fig 29.** Upregulated genes enrichment results for IL1 region

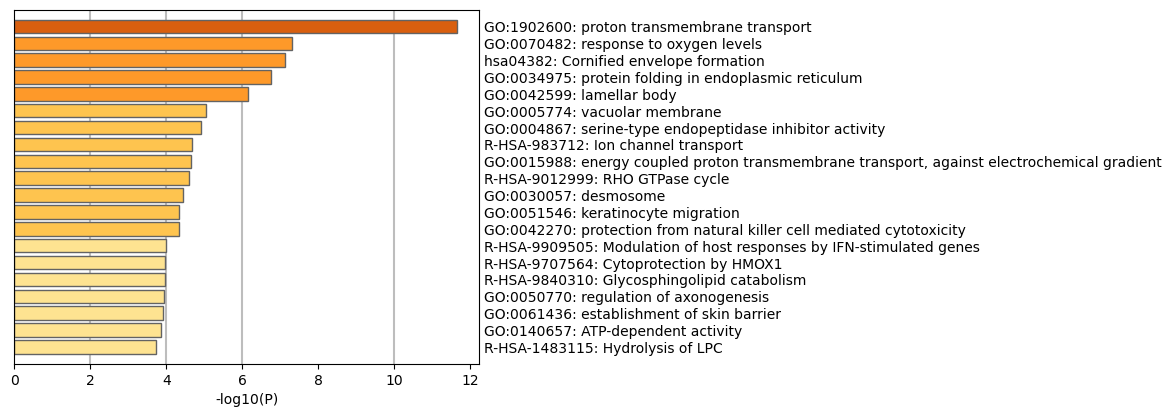

**Fig 30.** Upregulated genes enrichment results for IL2 region.

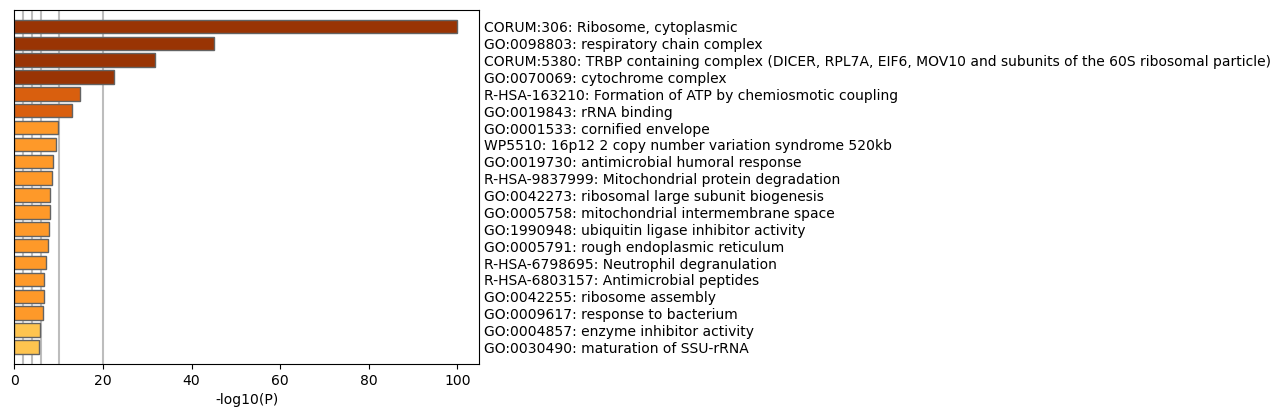

**Fig 31.** Upregulated genes enrichment results for IL3 region.

**KCs: KGF Cells**

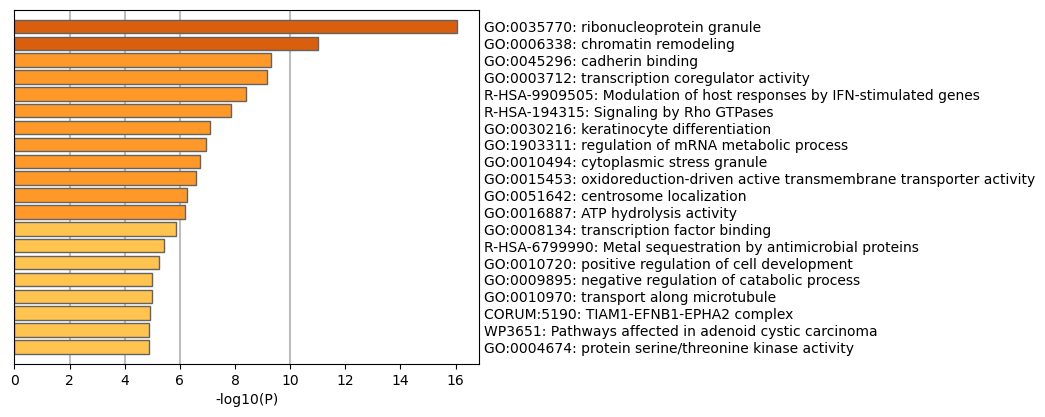

**Fig 32.** Downregulated genes enrichment results for K1 region.

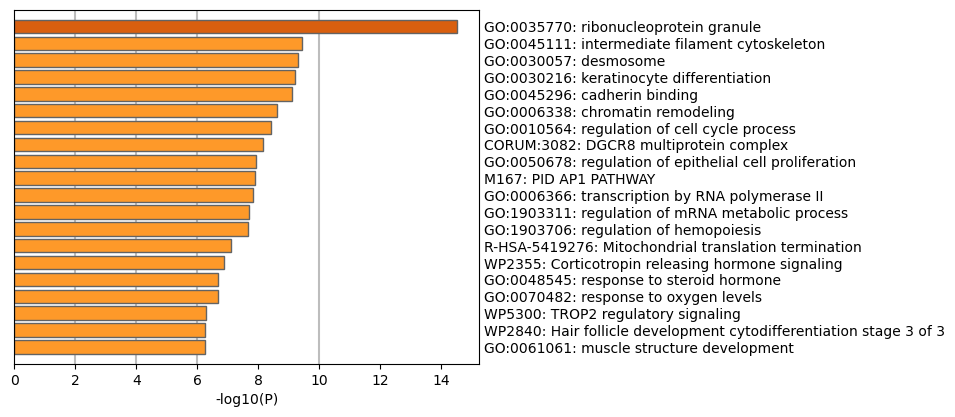

**Fig 33.** Downregulated genes enrichment results for K2 region.

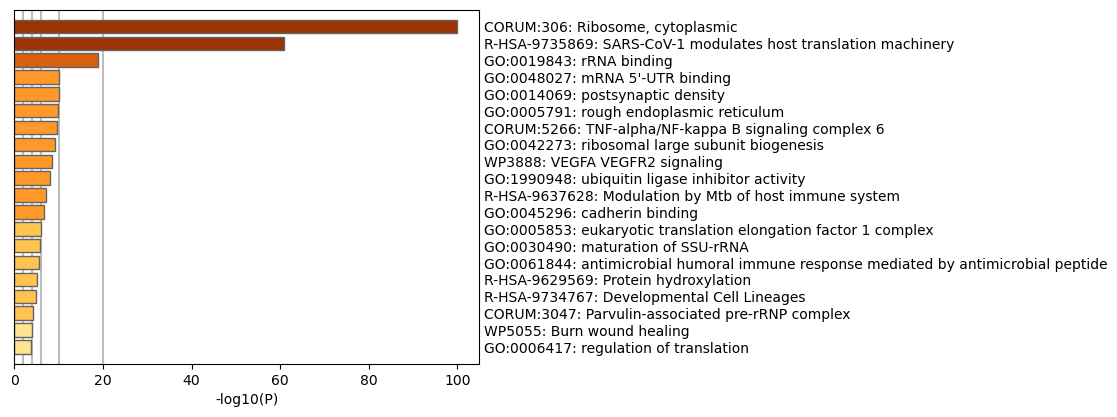

**Fig 34.** Downregulated genes enrichment results for K3 region.

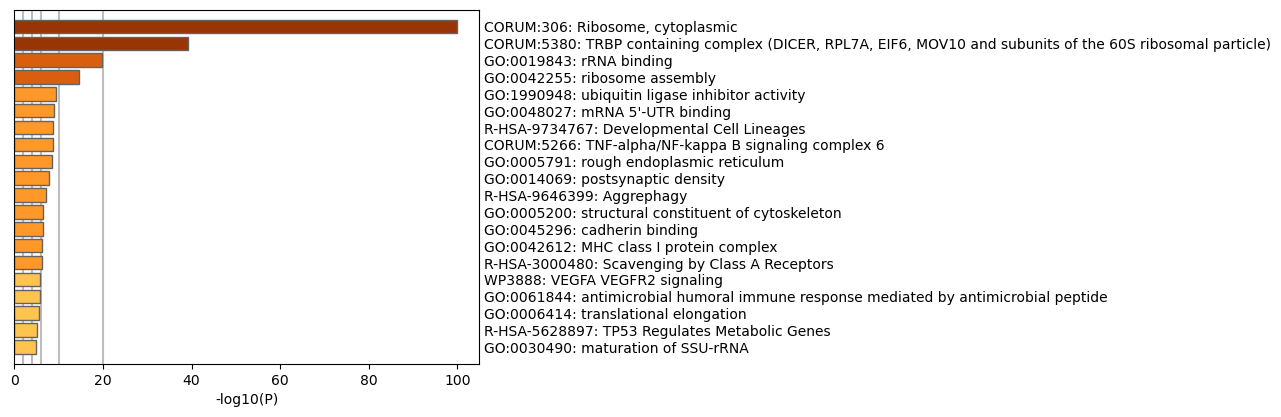

**Fig 35.** Upregulated genes enrichment results for K1 region.

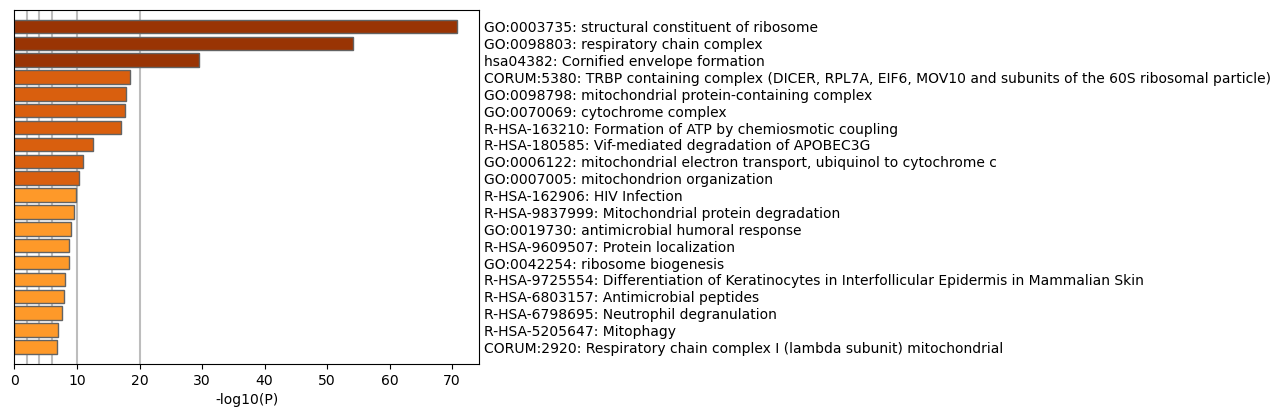

**Fig 36.** Upregulated genes enrichment results for K2 region.

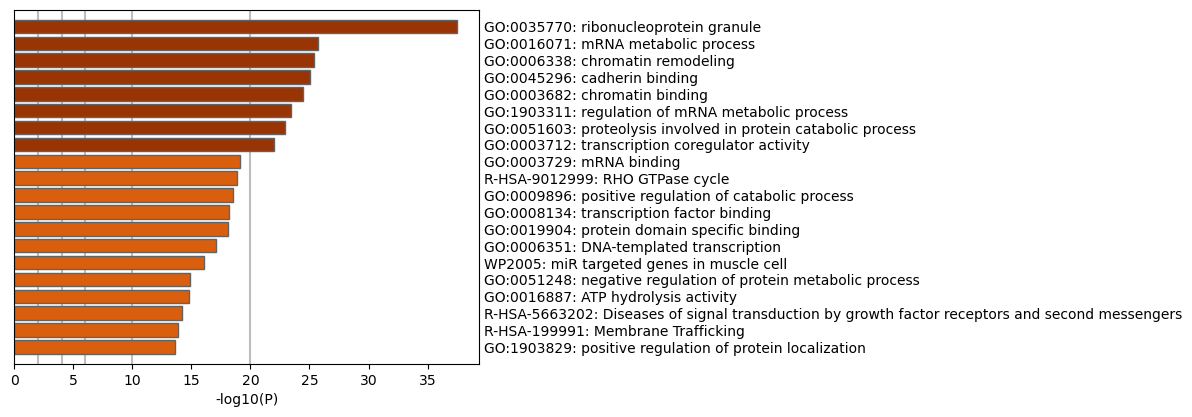

**Fig 37.** Upregulated genes enrichment results for K3 region.

**Neuron Cells**

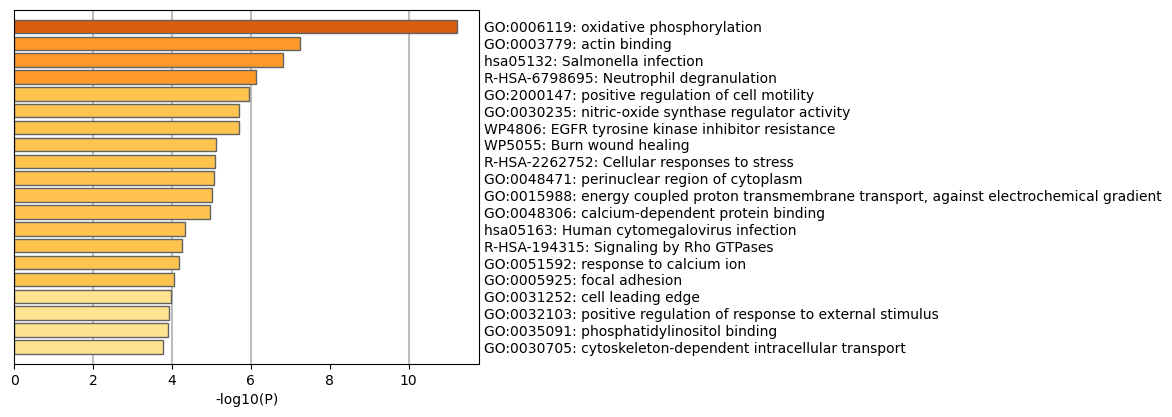

**Fig 38.** Downregulated genes enrichment results for N1 region.

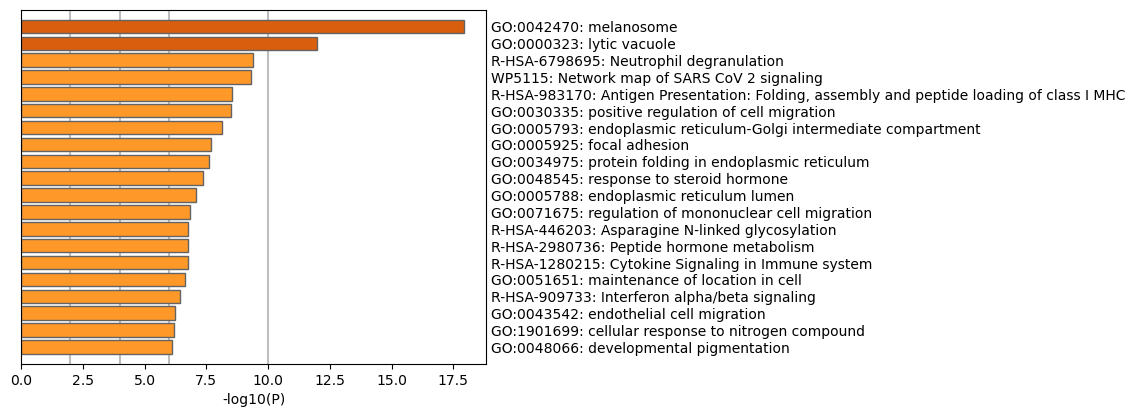

**Fig 39.** Downregulated genes enrichment results for N2 region.

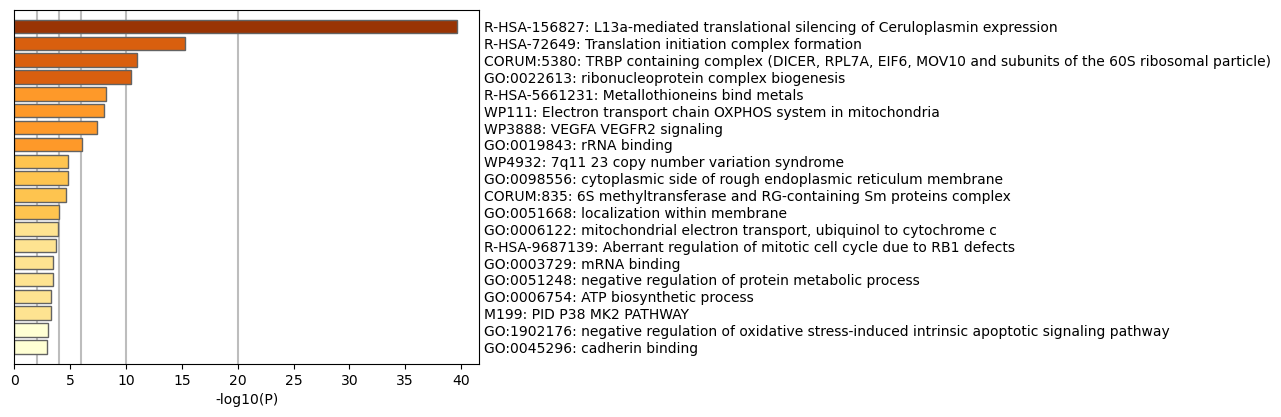

**Fig 40.** Downregulated genes enrichment results for N3 region.

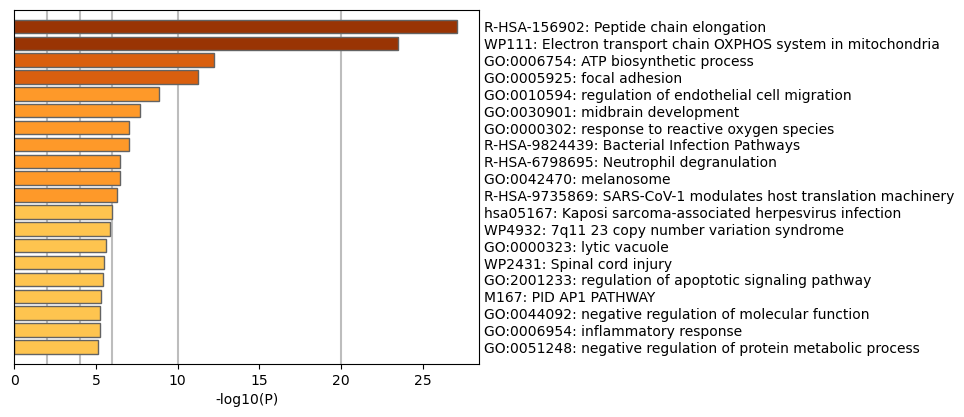

**Fig 41.** Upregulated genes enrichment results for N1 region.

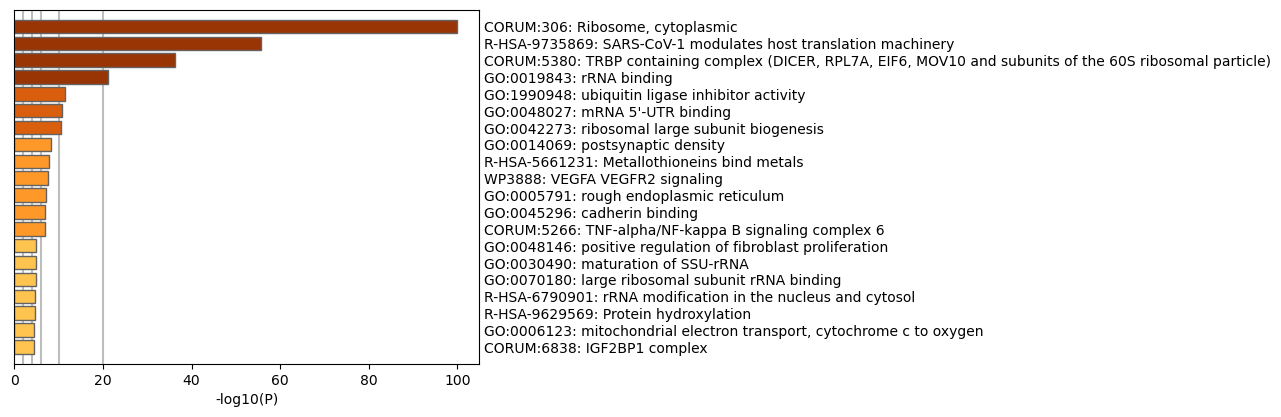

**Fig 42.** Upregulated genes enrichment results for N2 region.

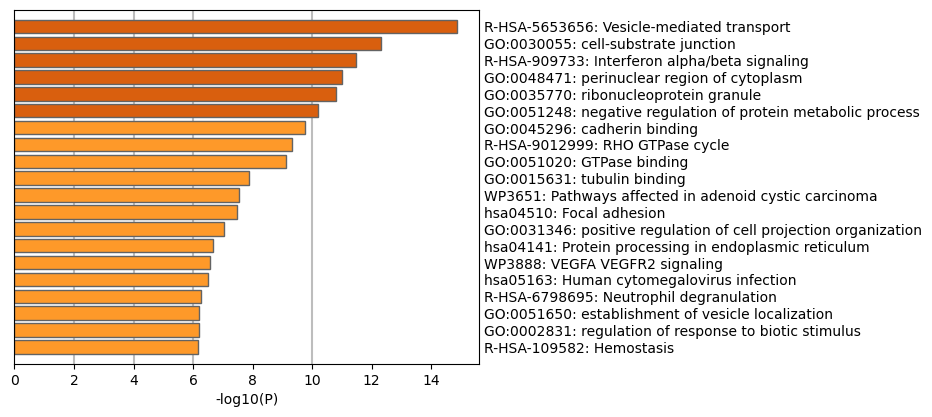

**Fig 43.** Upregulated genes enrichment results for N3 region.

**Group E (*disease-spesific signatures*) Enrichment Results**

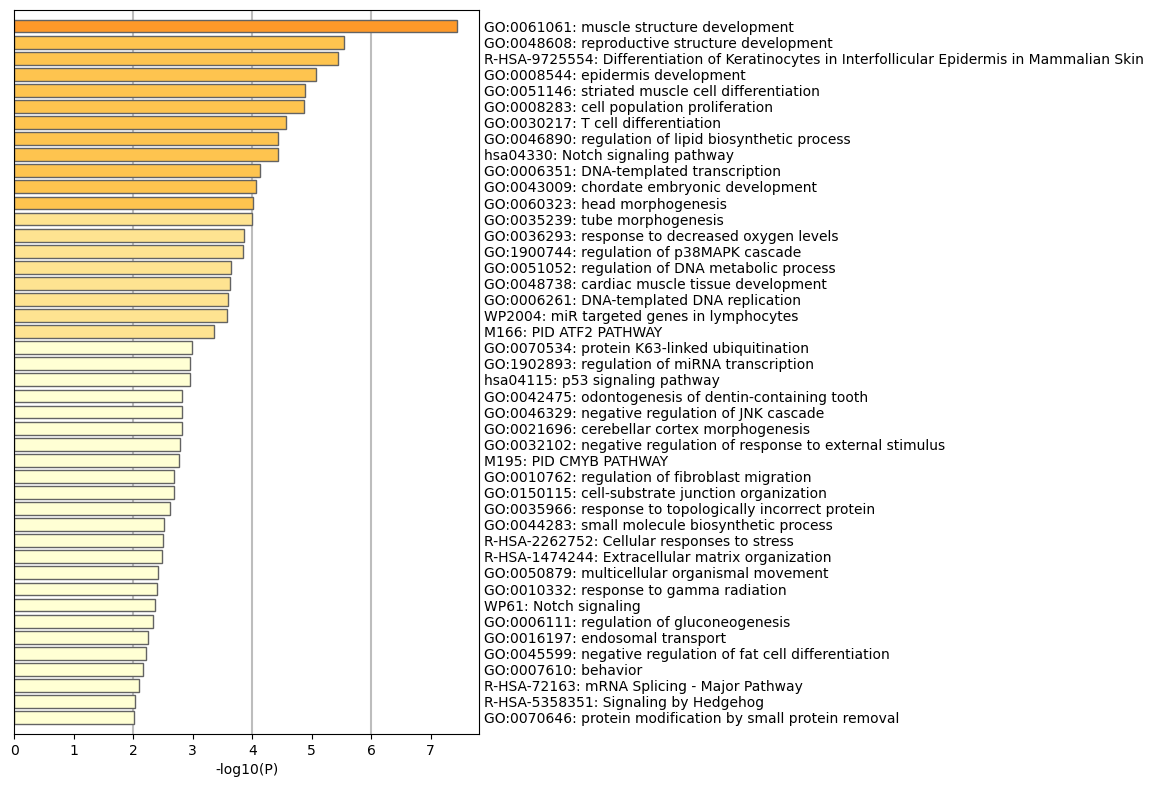

**Fig 44.** Upregulated genes enrichment results for Group E.

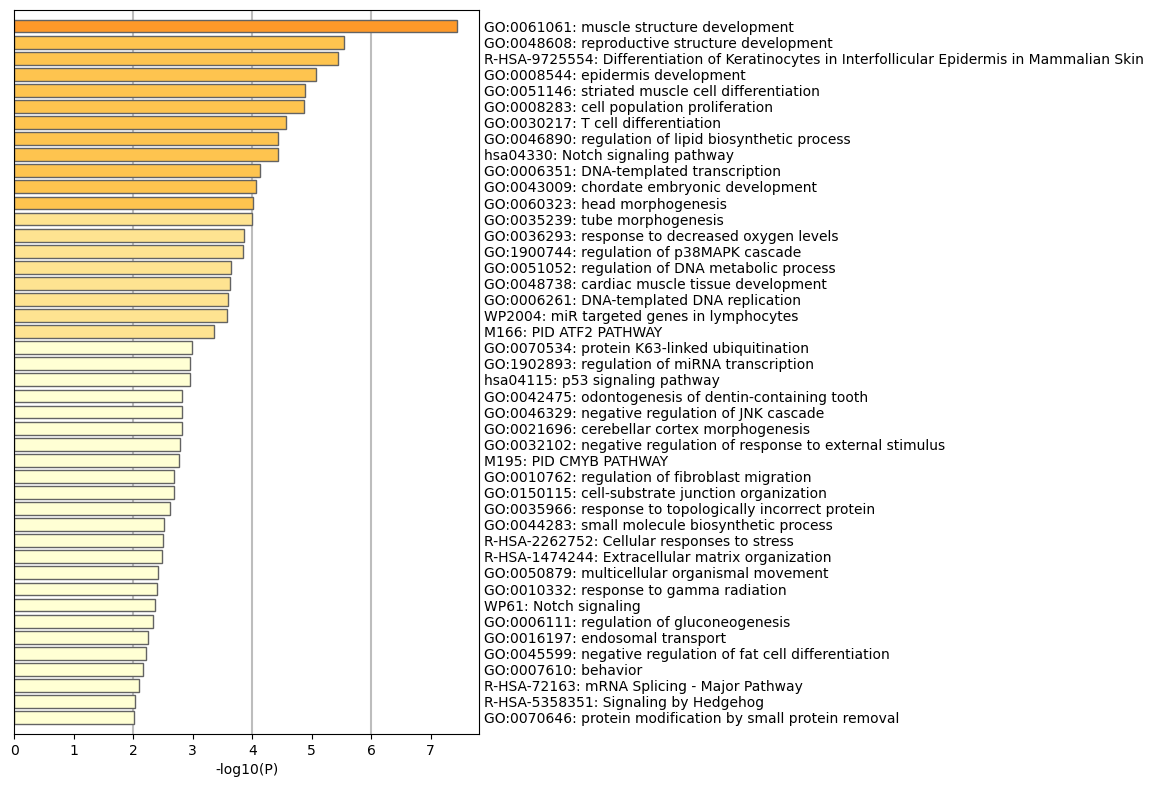

**Fig 45.** Downregulated genes enrichment results for Group E.
